# Control of spinal motor neuron terminal differentiation through sustained *Hoxc8* gene activity

**DOI:** 10.1101/2021.05.26.445841

**Authors:** Catarina Catela, Yifei Weng, Kailong Wen, Weidong Feng, Paschalis Kratsios

## Abstract

Spinal motor neurons (MNs) constitute cellular substrates for several movement disorders. Although their early development has received much attention, how spinal MNs become and remain terminally differentiated is poorly understood. Here, we determined the transcriptome of mouse brachial MNs at embryonic and postnatal stages. We found that genes encoding homeodomain (HOX, LIM) transcription factors (TFs), previously implicated in early MN development, continue to be expressed postnatally, suggesting later functions. To test this, we inactivated *Hoxc8* at successive stages of MN development. We found that *Hoxc8* is not only required to establish but also maintain expression of several MN terminal differentiation markers. Furthermore, we uncovered novel TFs with continuous MN expression, a *Hoxc8* dependency for maintained expression of Iroquois (*Irx*) homeodomain TFs, and a new role for *Irx2* in MN development. Our findings dovetail recent observations in *C. elegans* MNs, pointing toward an evolutionarily conserved role for Hox in neuronal terminal differentiation.

## INTRODUCTION

Motor neurons (MNs) represent the main output of our central nervous system. They control both voluntary and involuntary movement and are cellular substrates for several degenerative disorders(Arora and Khan, 2021). Due to their stereotypic cell body position, easily identifiable axons and highly precise synaptic connections with well-defined muscles, MNs are exceptionally well characterized in all major model systems. Extensive research over the past decades has focused on the early steps of MN development, thereby advancing our understanding of the molecular mechanisms controlling specification of progenitor cells and young post-mitotic MNs, as well as motor circuit assembly(Osseward and Pfaff, 2019, Philippidou and Dasen, 2013, Sagner and Briscoe, 2019). In the vertebrate spinal cord, progenitor cell specification critically depends on morphogenetic signals, whereas initial fate determination of post-mitotic MNs relies on combinatorial activity of different classes of transcription factors (TFs)(Dalla Torre di Sanguinetto et al., 2008, Jessell, 2000, Lee and Pfaff, 2001, Stifani, 2014). The focus in early development, however, has left poorly explored the molecular mechanisms that control the final steps of MN differentiation. Once MNs are born and specified, how do they acquire their terminal differentiation features, such as neurotransmitter phenotype, electrical and signaling properties? And perhaps most important, what are the mechanisms that ensure maintenance of such features throughout life?

The terminal differentiation features of every neuron type are determined by the expression of specific sets of proteins, such as neurotransmitter (NT) biosynthesis components, NT receptors, ion channels, neuropeptides, signaling molecules, trans-membrane receptors, and adhesion molecules (Hobert, 2008). The genes coding for these proteins (“terminal differentiation genes”) are continuously expressed from development through adulthood, thereby determining the functional and phenotypic properties of individual neuron types (Hobert, 2008, Hobert, 2011). Therefore, the challenge of understanding how MNs acquire and maintain their functional features lies in understanding how the expression of MN terminal differentiation genes is regulated over time. Importantly, defects in expression of such genes constitute one of the earliest molecular signs of MN disease (Nutini et al., 2011, Shibuya et al., 2011). However, the regulatory mechanisms that induce and maintain expression of terminal differentiation genes in spinal MNs are poorly defined. In part, this is due to: (a) a scarcity of temporally controlled gene inactivation studies that remove the activity of MN-expressed regulatory factors (e.g., TF, chromatin factor) at different life stages, and (b) a paucity of terminal differentiation markers for spinal MNs. Although recent RNA-Sequencing (RNA-Seq) studies have begun to address the latter (Blum et al., 2021, Delile et al., 2019, Alkaslasi et al., 2021), most genetic and molecular profiling studies on spinal MNs are not conducted in a longitudinal fashion, i.e., at embryonic and postnatal stages. Hence, how these cells become and remain terminally differentiated remains unclear.

To elucidate the molecular mechanisms that enable spinal MNs to acquire and maintain their terminal differentiation features, we took advantage of the orderly anatomical relationship between MN cell body location and muscle innervation, referred to as “topography” (Dasen and Jessell, 2009). In the spinal cord, this topographic relationship is mostly evident along the rostro-caudal axis, where MN populations located in different spinal cord domains (e.g., brachial, thoracic, lumbar, sacral) innervate different muscles. In this study, we solely focused on the brachial domain, where post-mitotic MNs are organized into two columns: (a) the lateral motor column [LMC] contains limb-innervating MNs necessary for reaching, grasping, and locomotion, and (b) the medial motor column [MMC] contains axial muscle-innervating MNs required for postural control(Philippidou and Dasen, 2013). Through a longitudinal RNA-Seq approach, we identified multiple terminal differentiation markers and novel TFs with continuous expression in embryonic and post-natal brachial MNs. Interestingly, we also found that several homeodomain TFs (HOX, LIM) that were previously implicated in the early steps of brachial MN development (e.g., initial specification, circuit assembly) (Philippidou and Dasen, 2013, Stifani, 2014) continue to be expressed in post-natal MNs. We therefore hypothesized that some of these TFs play additional roles in later steps of brachial MN development.

To test this hypothesis, we focused on Hox proteins because recent findings in the ventral nerve cord (equivalent to mouse spinal cord) of the nematode *C. elegans* identified Hox proteins as critical regulators of cholinergic MN terminal differentiation (Feng et al., 2020, Kratsios et al., 2017). Among the seven Hox genes retrieved from our RNA-Seq, *Hoxc8* is highly expressed both in embryonic and postnatal brachial MNs. A previous study showed that *Hoxc8* acts early to establish brachial MN connectivity (Catela et al., 2016). Here, we report a new role for *Hoxc8* in later stages of mouse MN development. By inactivating *Hoxc8* at successive developmental stages, we found that it is necessary for the establishment and maintenance of select terminal differentiation features of brachial MNs. Through an unbiased approach, we also identified new *Hoxc8* target genes in brachial MNs, including three members of the Iroquois family of homeodomain TFs *(Irx2, Irx5, Irx6).* Because mouse Hox genes are expressed both in the embryonic and postnatal hindbrain(Hutlet et al., 2016, Krumlauf, 2016), similar Hox-based mechanisms to the one described here may be widely used in the nervous system for the control of neuronal terminal differentiation.

## RESULTS

### Molecular profiling of mouse brachial MNs at embryonic and postnatal stages

We first sought to define the molecular profile of brachial MNs at embryonic and postnatal stages with the goal of identifying putative terminal differentiation markers for these cells. This longitudinal approach focused on post-mitotic MNs at embryonic day 12 (e12) and postnatal day 8 (p8). We chose e12 because: (i) spinal e12 MNs begin to acquire their terminal differentiation features, such as neurotransmitter phenotype (Martinez et al., 2012), and (ii) MN axons at e12 have exited the spinal cord (Catela et al., 2016). We chose p8 because: (i) this is several days after neuromuscular synapse formation (Gautam, 1996 #686), and (ii) pups at p8 become more active, indicating spinal MN functionality. To genetically label e12 MNs, we used the *Hb9::GFP* reporter mouse(Wichterle et al., 2002) (**Fig. 1A**). Due to low expression of *Hb9::GFP* at postnatal stages, we turned to an alternative labeling strategy and crossed *ChAT::IRES::Cre* mice(Rossi et al., 2011) with the *Ai9* Cre-responder line [Rosa26-CAG^promoter^-loxP-STOP-loxP-tdTomato] (Madisen et al., 2010). At p8, we observed fluorescent labeling of spinal MNs with tdTomato (**Fig. 1A**). Taking advantage of the topographic MN organization along the rostro-caudal axis, we followed a region-specific approach focused on the brachial region (segments C4-T1) that contains MNs of the MMC and LMC. Upon precise microdissection of this region (see Materials and Methods), we used fluorescence-activated cell sorting (FACS) to isolate GFP-labeled brachial MNs from e12 *Hb9::GFP* mice and tdTomato-labeled brachial MNs from p8 *ChAT::IRES::Cre;* Ai9 mice (**Fig. 1A**). Through RNA-Sequencing (RNA-Seq), we obtained and compared the molecular profile of these cells (see Materials and Methods). We identified differentially expressed transcripts (>4-fold, p < 0.05) in the e12 (3,715 transcripts) and p8 (3,209 transcripts) dataset (**Fig. 1B, Suppl. File 1**), suggesting gene expression profiles of embryonic and post-natal brachial MNs differ. We caution, however, that these differences could occur due to: (1) different levels of gene expression (see next Section), and (2) a fraction of the enriched transcripts is expressed in non-MNs because *Hb9* and *ChAT* are also expressed in small, non-overlapping interneuron populations (Wilson et al., 2005, Zagoraiou et al., 2009).

**Figure 1.**
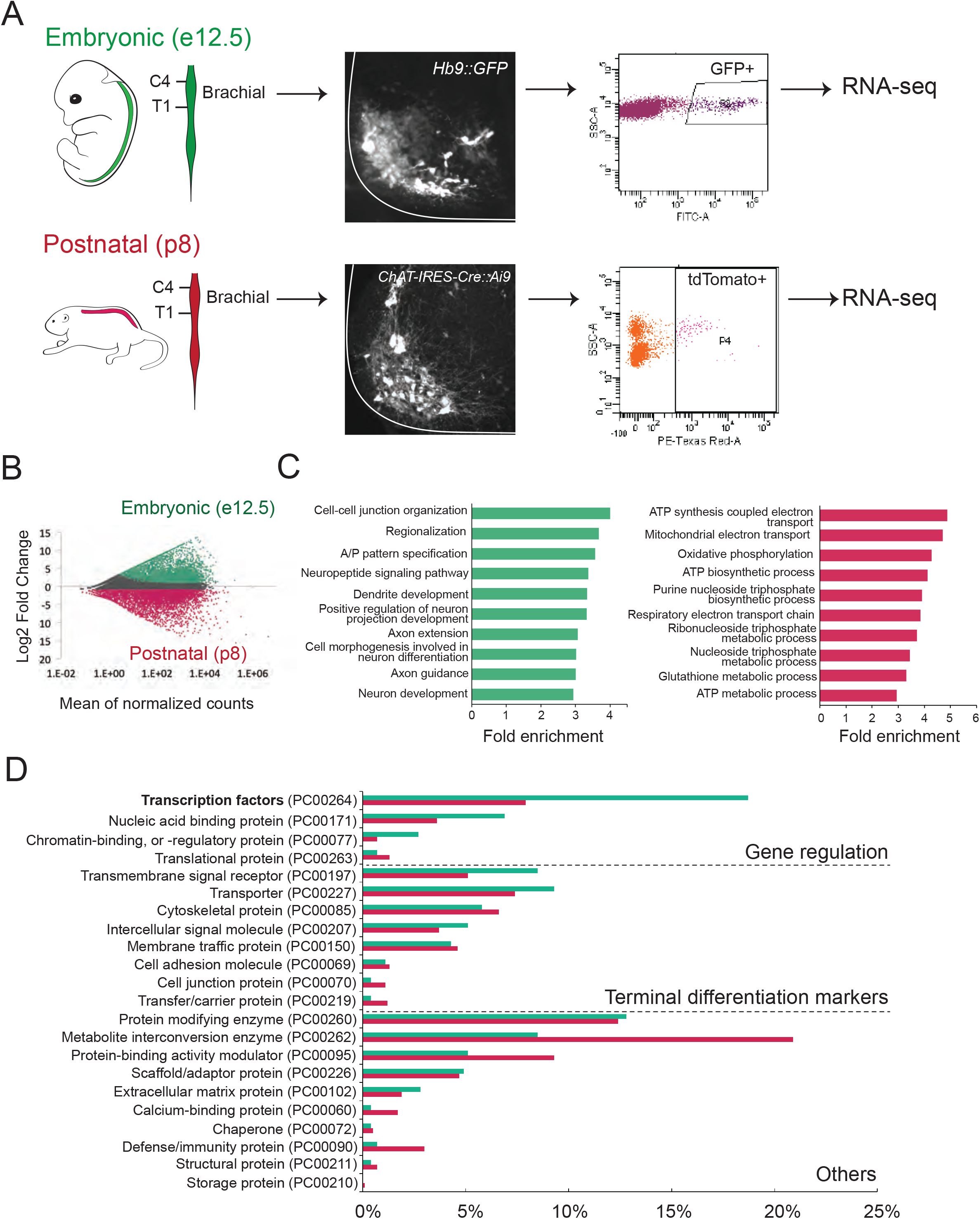
Molecular profiling of mouse brachial motor neurons at embryonic and postnatal stages. (A) Schematic representation of the workflow used in the comparison of embryonic and postnatal transcriptomes. The brachial domain (C4-T1) of *Hb9::Cre* (in green) and *ChAT-IRES-Cre:Ai9* (in red) mice was microdissected. Brachial GFP^+^ (at e12.5) and tdTomato^+^ (at p8) motor neurons were FACS-sorted and processed for RNA-sequencing. (B) MA plot of differentially expressed genes. Green and red dots represent individual genes that are significantly (p<0.05) expressed (4-fold and/or higher) in embryonic and postnatal MNs, respectively. (C) Graphs showing fold enrichment for genes involved in specific biological processes. (D) Gene Onthology (GO) analysis comparing protein class categories of highly expressed genes in embryonic (e12.5) and postnatal (p8) MNs. Green and red bars represent embryonic and postnatal genes, respectively.

Subsequent gene ontology (GO) analysis on proteins from embryonically enriched (e12) transcripts revealed an overrepresentation of molecules associated with neuronal development, such as regionalization, dendrite formation and axon guidance (**Fig. 1C, Suppl. File 2**). Notably, the most enriched class of proteins in the e12 dataset is TFs, many of which are known to control MN development (**Fig. 1D,** see next Section). On the other hand, GO analysis on proteins from postnatally enriched (p8) transcripts uncovered an overrepresentation of molecules associated with cell metabolism, such as ATP synthesis, oxidative phosphorylation and energy coupled proton transport (**Fig. 1C-D, Suppl. File 2**), perhaps indicative of the higher metabolic demands of p8 MNs compared to their embryonic (e12) counterparts.

To identify terminal differentiation markers with continuous expression in brachial MNs, we leveraged our e12 RNA-Seq dataset (**Fig. 1D, Suppl. File 1**). From this set, we arbitrarily selected 8 genes coding for neurotransmitter receptors, ion channels, and signaling molecules *(Slc10a4, Nrg1, Nyap2, Sncg, Ngfr, Glra2, Cldn1, Cacna1g)* and evaluated their expression at different life stages. Through RNA in situ hybridization (ISH), we found 6 genes *(Slc10a4, Nrg1, Nyap2, Sncg, Ngfr, Glra2)* with continuous expression in putative brachial MNs at embryonic (e12) and early postnatal (p8) stages (**Table 1, Suppl. Fig. 1**). Available RNA ISH data from the Allen Brain Atlas also confirmed their expression at p56 (**Table 1**). Although RNA ISH data lack single-cell resolution, the ventrolateral location of cells expressing these 6 genes strongly suggests they constitute terminal differentiation markers for brachial MNs.

**Table 1.**
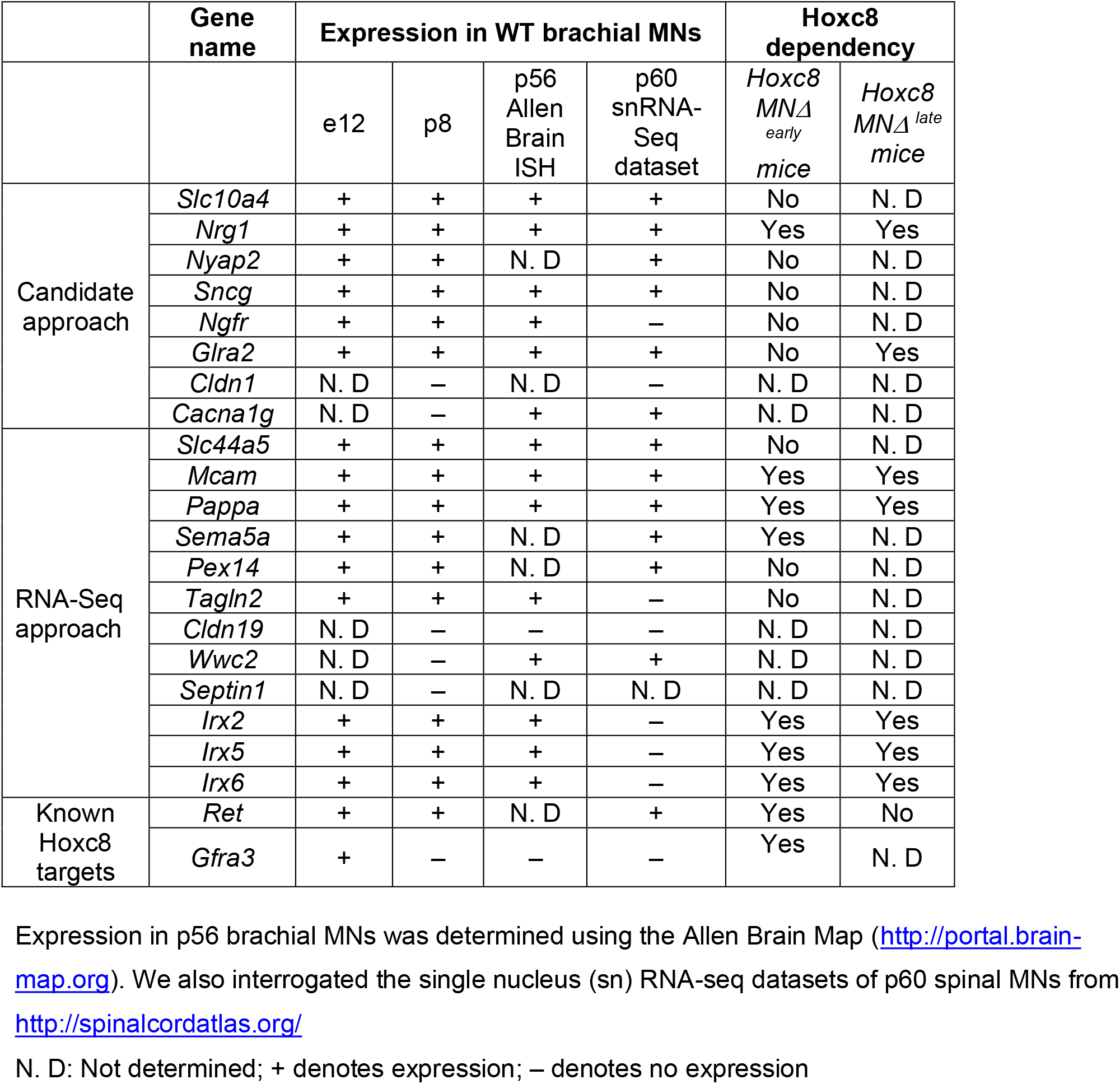
Summary of candidate and unbiased approaches to reveal Hoxc8 target genes in mouse brachial MNs.

### Developmental transcription factors continue to be expressed in spinal MNs at postnatal stages

Two simple, but not mutually exclusive mechanisms can be envisioned for the continuous expression of MN terminal differentiation genes. Their embryonic initiation and maintenance could be controlled by separate mechanisms involving distinct combinations of TFs solely dedicated to either initiation or maintenance. Alternatively, initiation and maintenance can be achieved through the activity of the same, continuously expressed TF (or combinations thereof). Recent invertebrate and vertebrate studies on various neuron types support the latter mechanism(Deneris and Hobert, 2014, Hobert and Kratsios, 2019). We therefore sought to identify TFs with continuous expression in mouse brachial MNs.

First, we examined whether TFs from our embryonic (e12) RNA-Seq dataset continue to be expressed at postnatal stages (**Fig. 1D**). We initially focused on 14 TFs from various families (e.g., LIM, Hox) with previously known embryonic expression and function in brachial MNs *(Ebf2, Islet1, Islet2, Hb9, Foxp1, Lhx3, Runx1, Hoxc4, Hoxa5, Hoxc5, Hoxa6, Hoxc6, Hoxa7, Hoxc8*)(Arber et al., 1999, Catela et al., 2019, Catela et al., 2016, Dasen et al., 2008, Ericson et al., 1992, Philippidou and Dasen, 2013, Sharma et al., 1998, Stifani et al., 2008, Thaler et al., 1999, Thaler et al., 2004, Thaler et al., 2002, Tsuchida et al., 1994). Through RNA ISH or antibody staining, we detected robust expression in brachial MNs at e12 for all 14 factors. Notably, 13 of these TFs continue to be expressed – albeit at lower levels – in brachial MNs at p8 (**Fig. 2A, Table 2**), suggesting these proteins – in addition to their known roles during early MN development – may exert other functions at later developmental and/or postnatal stages. Seven of these 13 proteins are TFs of the Hox family known to be expressed in brachial MNs at embryonic stages(Philippidou and Dasen, 2013), confirming regional specificity in the RNA-Seq approach (**Fig. 2A**). Moreover, our strategy is sensitive as it captured TFs with known expression in small populations of brachial MNs (e.g., MMC neurons), such as Ebf2 and Lhx3 (**Fig. 2A**)(Catela et al., 2019, Sharma et al., 1998).

**Figure 2.**
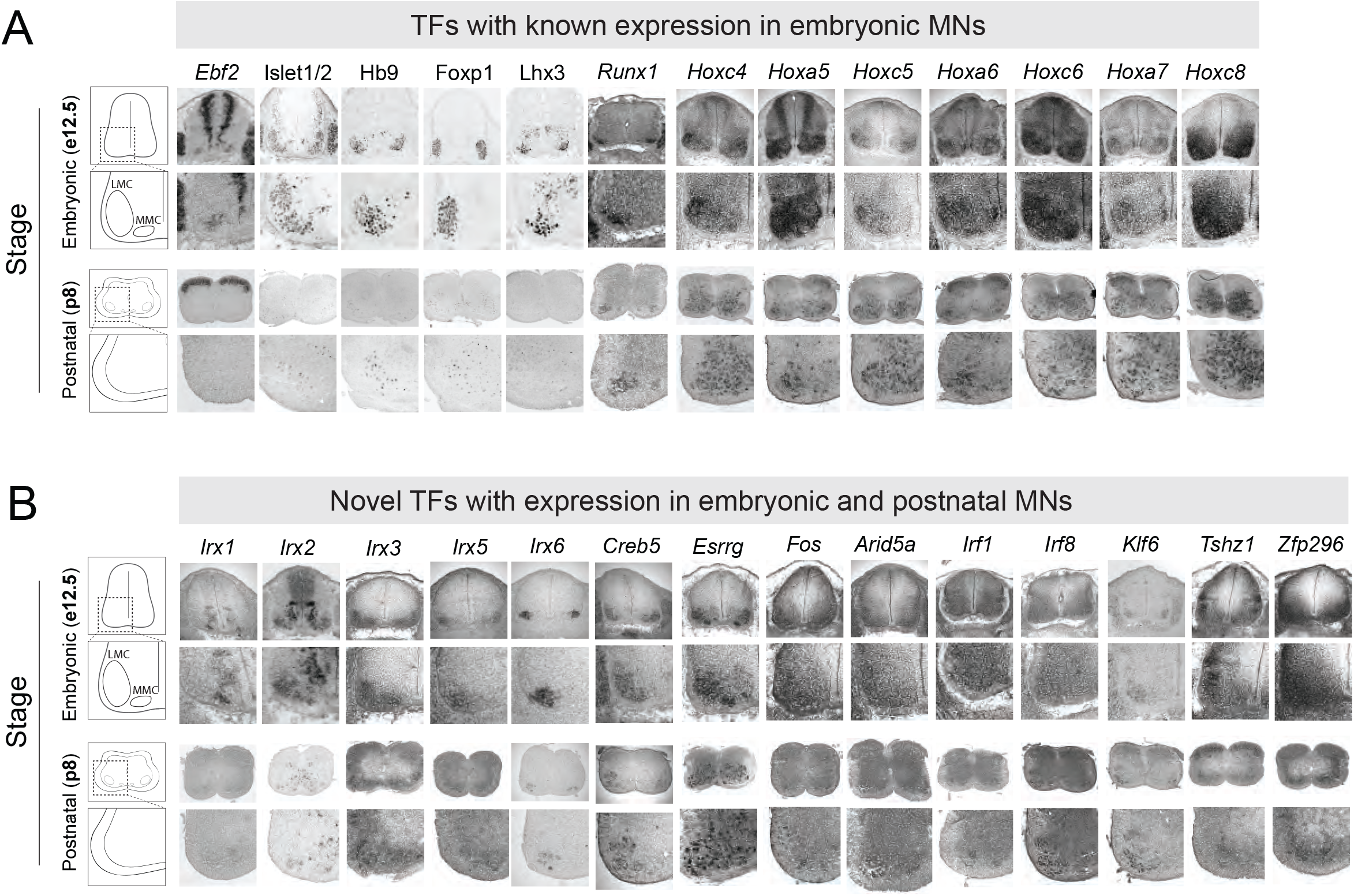
Known and novel transcription factors are continuously expressed in brachial motor neurons during embryonic and postnatal stages. (A) The expression of transcription factors with previously published roles in MN development was assessed in embryonic (e12.5) and postnatal (p8) spinal cords (N = 4) with RNA ISH *(Ebf2, Runx1, Hoxc4, Hoxa5, Hoxc5, Hoxa6, Hoxc6, Hoxa7, Hoxc8)* and immunohistochemistry (Islet1/2, Hb9, Lhx3, Foxp1). (B) The expression of novel transcription factors was assessed in embryonic (e12.5) and postnatal (p8) spinal cords with RNA ISH (N = 4).

**Table 2.**
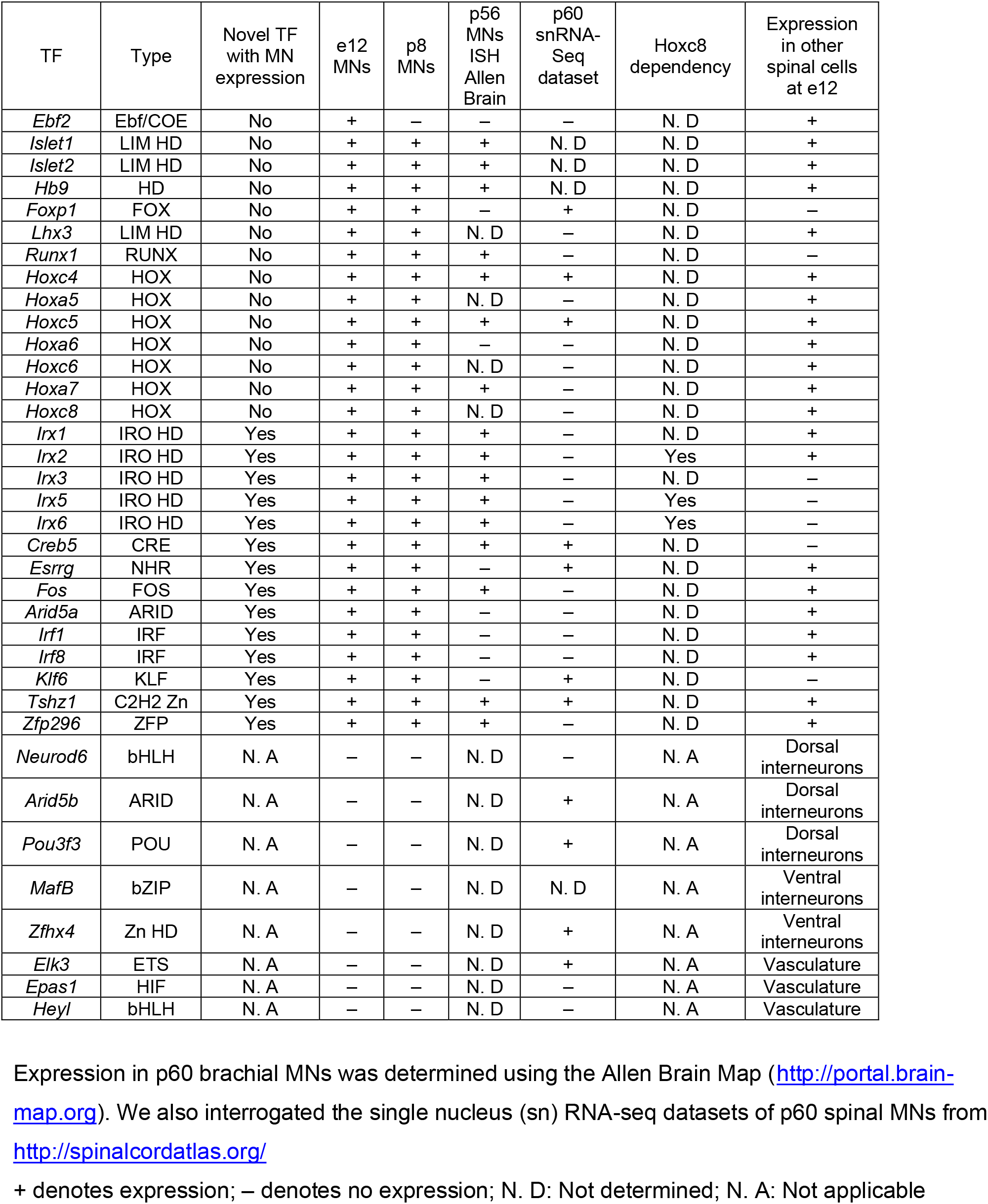
Validation of transcription factor expression in brachial MNs.

| TF | Type | Novel TF with MN expression | e12 MNs | p8 MNs | p56 MNs ISH Allen Brain | p60 snRNA-Seq dataset | Hoxc8 dependency | Expression in other spinal cells at e12 |
| --- | --- | --- | --- | --- | --- | --- | --- | --- |
| *Ebf2* | Ebf/COE | No | + | – | – | – | N. D | + |
| *Islet1* | LIM HD | No | + | + | + | N. D | N. D | + |
| *Islet2* | LIM HD | No | + | + | + | N. D | N. D | + |
| *Hb9* | HD | No | + | + | + | N. D | N. D | + |
| *Foxp1* | FOX | No | + | + | – | + | N. D | – |
| *Lhx3* | LIM HD | No | + | + | N. D | – | N. D | + |
| *Runx1* | RUNX | No | + | + | + | – | N. D | – |
| *Hoxc4* | HOX | No | + | + | + | + | N. D | + |
| *Hoxa5* | HOX | No | + | + | N. D | – | N. D | + |
| *Hoxc5* | HOX | No | + | + | + | + | N. D | + |
| *Hoxa6* | HOX | No | + | + | – | – | N. D | + |
| *Hoxc6* | HOX | No | + | + | N. D | – | N. D | + |
| *Hoxa7* | HOX | No | + | + | + | – | N. D | + |
| *Hoxc8* | HOX | No | + | + | N. D | – | N. D | + |
| *Irx1* | IRO HD | Yes | + | + | + | – | N. D | + |
| *Irx2* | IRO HD | Yes | + | + | + | – | Yes | + |
| *Irx3* | IRO HD | Yes | + | + | + | – | N. D | – |
| *Irx5* | IRO HD | Yes | + | + | + | – | Yes | – |
| *Irx6* | IRO HD | Yes | + | + | + | – | Yes | – |
| *Creb5* | CRE | Yes | + | + | + | + | N. D | – |
| *Esrrg* | NHR | Yes | + | + | – | + | N. D | + |
| *Fos* | FOS | Yes | + | + | + | – | N. D | + |
| *Arid5a* | ARID | Yes | + | + | – | – | N. D | + |
| *Irf1* | IRF | Yes | + | + | – | – | N. D | + |
| *Irf8* | IRF | Yes | + | + | – | – | N. D | + |
| *Klf6* | KLF | Yes | + | + | – | + | N. D | – |
| *Tshz1* | C2H2 Zn | Yes | + | + | + | + | N. D | + |
| *Zfp296* | ZFP | Yes | + | + | + | – | N. D | + |
| *Neurod6* | bHLH | N. A | – | – | N. D | – | N. A | Dorsal interneurons |
| *Arid5b* | ARID | N. A | – | – | N. D | + | N. A | Dorsal interneurons |
| *Pou3f3* | POU | N. A | – | – | N. D | + | N. A | Dorsal interneurons |
| *MafB* | bZIP | N. A | – | – | N. D | N. D | N. A | Ventral interneurons |
| *Zfhx4* | Zn HD | N. A | – | – | N. D | + | N. A | Ventral interneurons |
| *Elk3* | ETS | N. A | – | – | N. D | + | N. A | Vasculature |
| *Epas1* | HIF | N. A | – | – | N. D | – | N. A | Vasculature |
| *Heyl* | bHLH | N. A | – | – | N. D | – | N. A | Vasculature |

Expression in p60 brachial MNs was determined using the Allen Brain Map ([http://portal.brain-](http://portal.brain-map.org) [map.org](http://portal.brain-map.org)). We also interrogated the single nucleus (sn) RNA-seq datasets of p60 spinal MNs from http://spinalcordatlas.org/ + denotes expression; – denotes no expression; N. D: Not determined; N. A: Not applicable

We next sought to identify novel TFs with maintained expression in brachial MNs. We arbitrarily selected 22 genes from different TF families (15 TFs from the e12 dataset [Irx1, Irx2, Irx3, Irx5, Irx6, Creb5, Esrrg, Neurod6, Arid5b, Pou3f3, MafB, Zfhx4, Elk3, Epas1, Heyl] and 7 TFs from the p8 dataset [Fos, Arid5a, Irf1, Irf8, Klf6, Tshz1, Zfp296]). We detected persistent expression for 14 of these TFs in the embryonic (e12) and early postnatal (p8) brachial spinal cord. Expression was evident at the ventrolateral region – a location populated by MNs (**Fig. 2B, Table 2**).

In conclusion, the expression of 13 TFs, with known roles in early MN development (e.g., cell specification, motor circuit assembly), is persistent at early postnatal stages (p8). Moreover, we identified 14 novel TFs from different families with expression in embryonic and postnatal (p8) brachial MNs (**Fig. 2B, Table 2**). The continuous expression of all these factors suggests that they may exert various functions in post-mitotic MNs at different life stages. Consistent with this notion, some of these TFs are also expressed at later postnatal (p56, p60) stages in brachial MNs (**Table 2**).

### Hoxc8 controls expression of several terminal differentiation genes in e12 brachial MNs

In mice, Hox genes play critical roles during the early steps of spinal cord development, such as MN specification and circuit assembly(Dasen et al., 2008, Dasen et al., 2003, Dasen et al., 2005, Philippidou and Dasen, 2013). We found that several Hox genes are continuously expressed – from embryonic to postnatal stages – in brachial MNs (**Fig. 2A**), but their function during later stages of MN development is largely unknown. This pattern of continuous Hox gene expression is reminiscent of recent observations in *C. elegans* nerve cord MNs(Feng et al., 2020, Kratsios et al., 2017). Importantly, *C. elegans* Hox genes are required not only to establish but also maintain at later stages the expression of multiple terminal differentiation genes (e.g., NT receptors, ion channels, signaling molecules) in nerve cord MNs(Feng et al., 2020).

Motivated by these findings in invertebrates, we sought to test the hypothesis that, in mice, Hox proteins control expression of terminal differentiation genes in spinal MNs. We focused on Hoxc8 because it is expressed in the majority of brachial MNs (segments C6-T2) (**Fig. 2A**) (Catela et al., 2016). Hoxc8 is not required for the overall organization of brachial MNs into columns, but – during early development (e12) – it controls forelimb muscle innervation by regulating *Gfrα3* and *Ret* expression in brachial MNs(Catela et al., 2016). However, whether Hoxc8 is involved in additional processes, such as the control of MN terminal differentiation, remains unclear.

To test this, we removed *Hoxc8* gene activity in brachial MNs. Because *Hoxc8* is also expressed in other spinal neurons(Baek et al., 2019, Shin et al., 2020), we crossed *Hoxc8^fl/fl^* mice to *Olig2::Cre* mice that enable *Cre* recombinase expression specifically in MN progenitors (**Fig. 3A**)(Zawadzka et al., 2010). This genetic strategy effectively removed Hoxc8 protein from post-mitotic brachial MNs by e12 (**Fig. 3B**). Because e12 is an early stage of MN differentiation (post-mitotic MNs are generated between e9 – e11)(Sims and Vaughn, 1979), we will refer to the *Olig2::Cre; Hoxc8^fl/fl^* mice as *Hoxc8 MNΔ^early^*. Of note, the total number of brachial MNs is unaffected in these animals at e12 (**Fig. 3C**).

**Figure 3.**
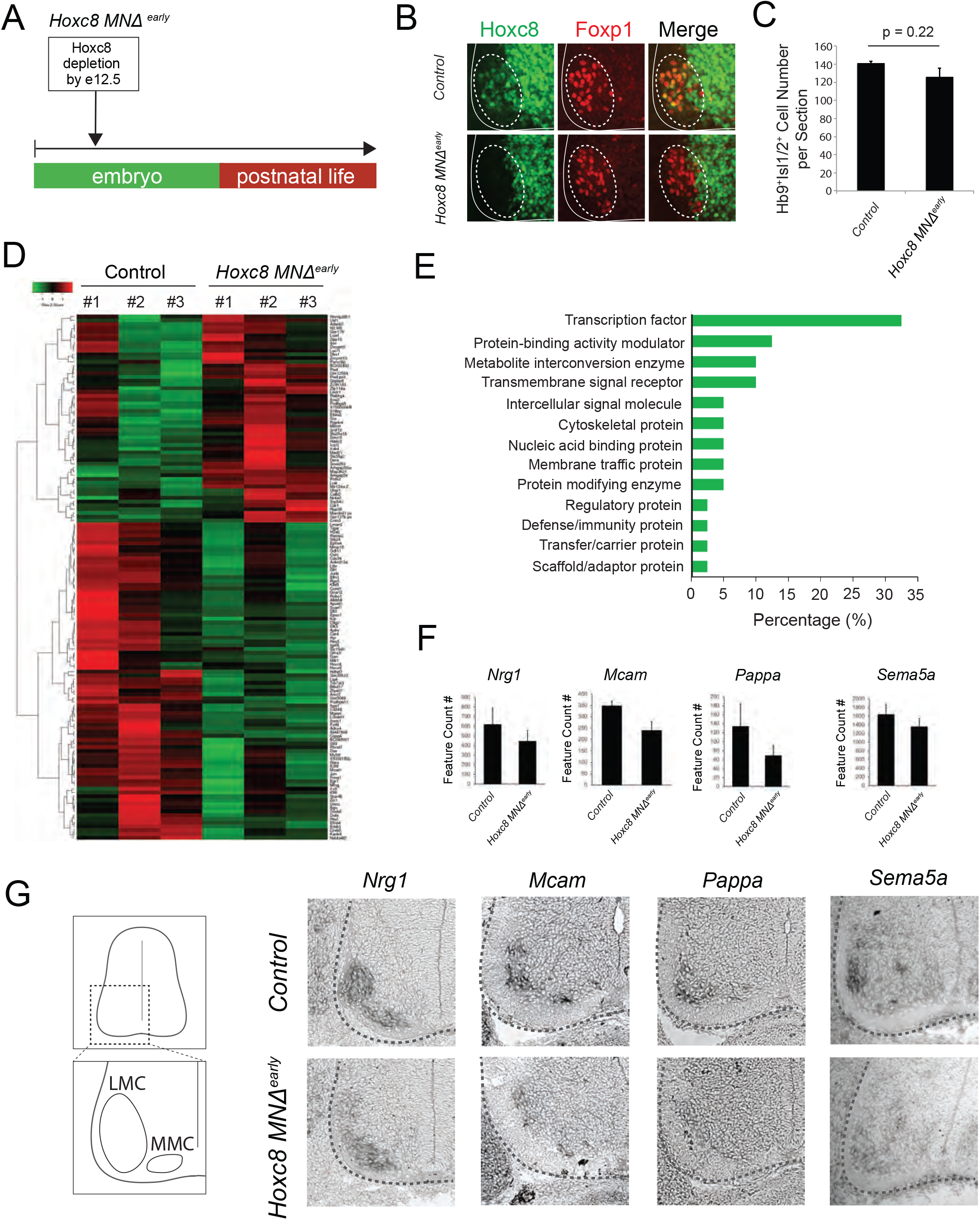
Early *Hoxc8* gene inactivation in brachial motor neurons affects the expression of terminal differentiation genes. (A) Diagram illustrating genetic approach for *Hoxc8* gene inactivation during early MN development. *Hoxc8* conditional mice were crossed with the *Olig2-Cre* mouse line *(Hoxc8 MNΔ ^early^*). (B) Immunohistochemistry showing that Hoxc8 protein (green) cannot be detected in Foxp1^+^ motor neurons (red) of *Hoxc8 MNΔ^early^* spinal cords at e12. (C) Quantification of Hb9^+^Isl1/2^+^ motor neurons in e12.5 brachial spinal cords of *Hoxc8 MNΔ^early^* and control embryos (N = 4). (D) Heatmap showing up-regulated and down-regulated genes detected by RNA-seq in control *(Hoxc8^fl/fl^* and *Hoxc8 MNΔ^early^* e12.5 motor neurons. Green and red colors respectively represent lower and higher gene expression levels. (E) Graphical percentage (%) representation of protein classes of the down-regulated genes in *Hoxc8 MNΔ^early^* spinal cords. (F) Graphical representation of the reduced count numbers of *Nrg1, Mcam, Pappa* and *Sema5a* mRNAs in *Hoxc8 MNΔ^early^* and control *(Hoxc8^fl/fl^)* spinal cords at e12 (N = 3). (G) RNA ISH showing downregulation of *Nrg1, Mcam, Pappa* and *Sema5a* mRNAs in brachial MNs of e12.5 *Hoxc8 MNΔ^early^* spinal cords (N = 4).

To test whether Hoxc8 controls expression of terminal differentiation genes, we initially followed a candidate approach. At e12, spinal MNs begin to acquire their terminal differentiation features, evident by the induction of genes coding for acetylcholine (ACh) biosynthesis proteins *(VAChT/Slc18a3, ChT1/Slc5a7)* (Martinez et al., 2012). Consistently, *VAChT/Slc18a3* and *ChT1/Slc5a7* transcripts were captured in our e12 RNA-Seq dataset (**Fig. 1D**). However, *VAChT/Slc18a3* and *ChT1/Slc5a7* expression was not affected in brachial MNs of *Hoxc8 MNΔ^eary^* mice (**Suppl. Fig. 1**). Next, we tested the six newly identified terminal differentiation markers *(Slc10a4, Nrg1, Nyap2, Sncg, Ngfr, Glra2)* summarized in **Table 1**. We found that expression of *Neuregulin 1 (Nrg1),* a molecule required for neuromuscular synapse maintenance and neurotransmission(Mei and Xiong, 2008, Wolpowitz et al., 2000), is reduced in e12 brachial MNs of *Hoxc8 MNΔ^early^* mice (**Fig. 3F-G**). However, the expression of the remaining five genes was unaffected in these animals (**Suppl. Fig. 1**), prompting us to devise an unbiased strategy.

We performed RNA-Seq on FACS-sorted brachial MNs from *Hoxc8 MNΔ^eary^; Hb9::GFP* and control mice at e12 (see Materials and Methods). We found dozens of significantly (p<0.05) up-regulated (55) and down-regulated (84) transcripts in MNs lacking *Hoxc8* (**Fig. 3D**). To test the hypothesis of *Hoxc8* being necessary for expression of MN terminal differentiation genes, we specifically focused on the list of 84 down-regulated transcripts, which included two known Hoxc8 target genes *(Ret, Gfrα3*)(Catela et al., 2016) and Hoxc8 itself (**Suppl. File 3**). GO analysis (see Materials and Methods) on these 84 transcripts identified several putative Hoxc8 target genes encoding proteins from various classes (**Fig. 3E, Suppl. File 3**). We focused on ion channels, transmembrane proteins, cell adhesion and signaling molecules, as these constitute putative terminal differentiation markers (Hobert, 2008, Hobert, 2011). We selected 9 genes *(Slc44a5, Mcam, Pappa, Sema5a, Pex14, Tagln2, Cldn19, Wwc2, Septin1)* and evaluated their expression with RNA ISH in brachial MNs at different stages. Five of these genes *(Slc44a5, Mcam, Pappa, Pex14, Tagln2)* are continuously expressed in brachial MNs at embryonic and postnatal stages (**Table 1, Suppl. Fig. 1**). Importantly, we found that expression of *Mcam,* a transmembrane cell adhesion molecule of the Immunoglobulin superfamily (Gu et al., 2015, Taira et al., 2004), and *Pappa,* a secreted molecule involved in skeletal muscle development(Rehage et al., 2007), is reduced at e12 in brachial MNs of *Hoxc8 MNΔ^early^* mice (**Fig. 3F-G, Suppl. Fig. 1**). In addition, we observed that *Sema5a* is expressed in embryonic (e12) but not postnatal brachial MNs, and this embryonic expression depends on Hoxc8 (**Table 1, Fig. 3F-G**). Because *Sema5a* encodes a transmembrane protein of the semaphorin protein family involved in axon guidance(Duan et al., 2014, Hilario et al., 2009, Lin et al., 2009), its dependency on *Hoxc8* could, at least partially, account for the previously reported motor neuron axonal defects of *Hoxc8 MNΔ^early^* mice (Catela et al., 2016).

Altogether, this analysis identified eleven terminal differentiation genes with continuous expression in brachial MNs *(Slc10a4, Nrg1, Nyap2, Sncg, Ngfr, Glra2, Slc44a5, Mcam, Pappa, Pex14, Tagln2),* three of which *(Nrg1, Mcam, Pappa)* constitute Hoxc8 targets (**Table 1**). Although additional, yet-to-be identified TFs must regulate the remaining eight genes, our findings do suggest Hoxc8 is involved in MN terminal differentiation. This new role for Hox in vertebrate MN development is consistent with recent studies in the *C. elegans* nerve cord, where Hox genes also control MN terminal differentiation (Feng et al., 2020, Kratsios et al., 2017).

### Hoxc8 is required to maintain expression of terminal differentiation genes in brachial MNs

Our analysis of *Hoxc8 MNΔ^early^* mice at e12 suggests Hoxc8 controls the early expression of select terminal differentiation genes *(Nrg1, Mcam, Pappa)* in brachial MNs. However, the persistent expression of *Hoxc8* both in embryonic and early postnatal MNs raises the intriguing possibility of a continuous requirement (**Fig. 2A**). Is *Hoxc8* required at later stages to maintain expression of terminal differentiation genes and thereby ensure the functionality of brachial MNs?

To address this, we crossed the *Hoxc8^fl/fl^* mice with the *ChAT::IRES::Cre* mouse line, which enables efficient gene inactivation in spinal MNs at later developmental stages when compared to *Hoxc8 MNΔ^early^* mice (**Fig. 4A**). Indeed, expression of *Hoxc8* mRNA persists at least until e14.5 in brachial MNs of *ChAT::IRES::Cre; Hoxc8^fl/fl^* mice (**Fig. 4B**). Given that post-mitotic MNs are generated between e9.5 and e11.5(Sims and Vaughn, 1979), this genetic strategy preserves *Hoxc8* expression in MNs at least for 3 days after their generation. Although we were unable to pinpoint the exact developmental stage (after e14.5) for onset of *Hoxc8* depletion in *ChAT::IRES::Cre; Hoxc8^fl/fl^* mice, we did witness loss of *Hoxc8* expression at a later stage (p8) in the ventrolateral region of the brachial spinal cord (location of MN cell bodies) (**Fig. 4B**). We will therefore refer to these animals as *Hoxc8 MNΔ^late^* because *Hoxc8* depletion in MNs occurs later compared to *Hoxc8 MNΔ^early^* mice (**Fig. 4A**). Interestingly, expression of the same terminal differentiation genes *(Nrg1, Mcam, Pappa)* we found affected in *Hoxc8 MNΔ^early^* mice is also reduced in brachial MNs of *Hoxc8 MNΔ^late^* mice at p8 (**Fig. 4C**). This reduction is not due to secondary events affecting MN generation or survival because similar numbers of brachial MNs were observed in control and *Hoxc8 MNΔ^late^* spinal cords at p8 (**Fig. 4D**). Taken together, our findings on *Hoxc8 MNΔ^early^* and *Hoxc8 MNΔ^late^* mice strongly suggest a continuous requirement – *Hoxc8* is required to establish and maintain at later developmental stages the expression of several terminal differentiation genes in brachial MNs (**Fig. 4F**).

**Figure 4.**
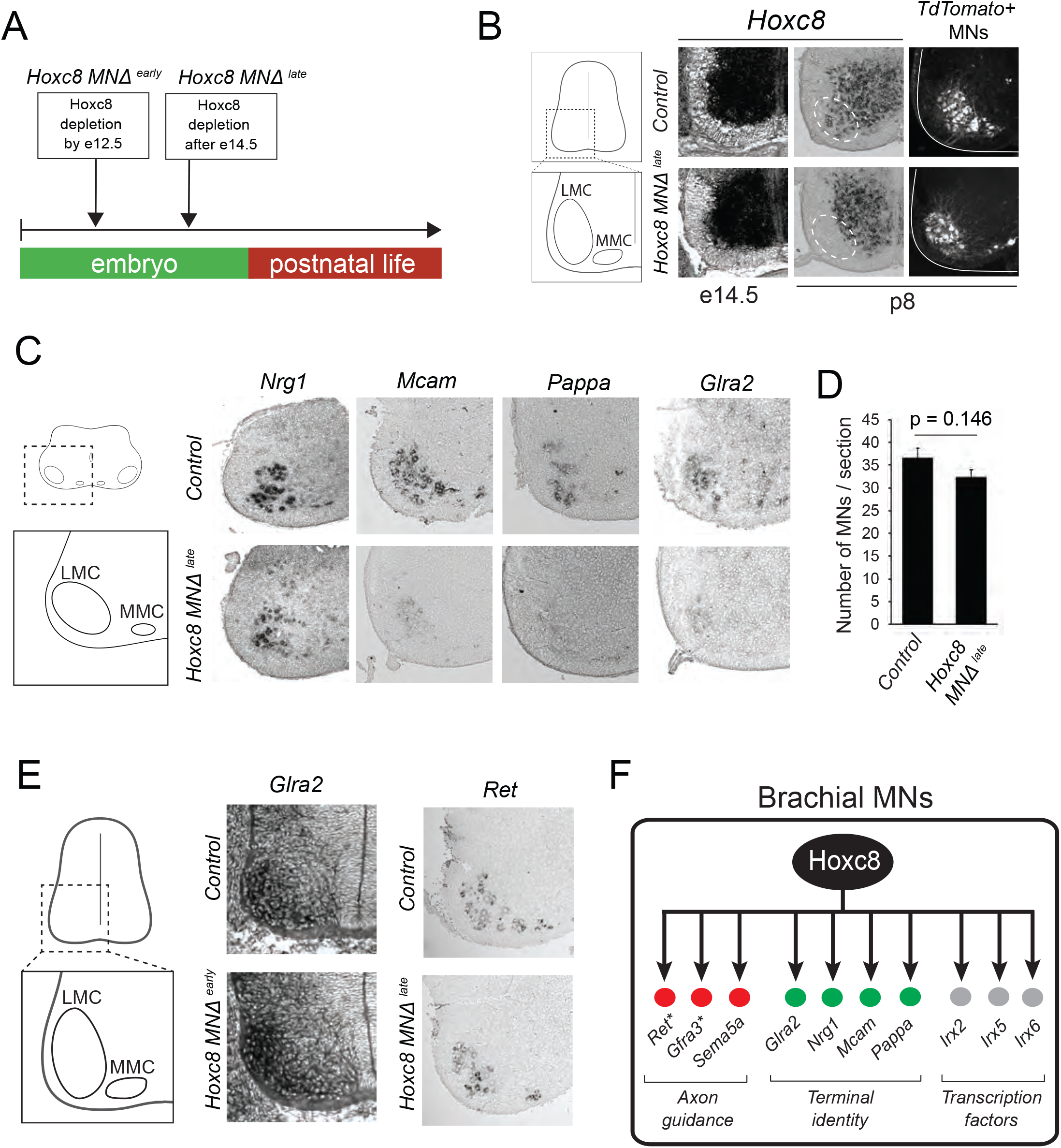
Late *Hoxc8* gene inactivation in brachial motor neurons affects expression of terminal differentiation genes. (A) Diagram illustrating genetic approach for *Hoxc8* gene inactivation during late MN development. *Hoxc8* conditional mice were crossed with the *ChAT-IRES-Cre* mouse line *(Hoxc8 MNΔ^late^*). (B) RNA ISH showing that *Hoxc8* mRNA is still present at e14.5 in *Hoxc8 MNΔ^late^* spinal cords, however, it cannot be detected at postnatal day 8 (p8) spinal cords of the same genotype (N=4). Representative images of td Tomato-labelled brachial MNs from *Hoxc8 ^fl/fl^; Ai9* (control) and *Hoxc8 MNΔ^late^; Ai9* spinal cords at p8. (C) RNA ISH showing reduced expression of *Pappa, Mcam, Glra2* and *Nrg1* in *Hoxc8 MNΔ^late^* spinal cords at p8 (N = 4). (D) Quantification of ChAT^+^ motor neurons in p8 control and *Hoxc8 MNΔ^late^* and spinal cords (N = 4). (E) RNA ISH showing that *Glra2* expression is comparable between control and *Hoxc8 MNΔ^early^* spinal cords at e12. Similarly, *Ret* expression is comparable between control and *Hoxc8 MNΔ^late^* spinal cords at p8 (N = 4). (F) Schematic summarizing *Hoxc8* target genes in brachial MNs. With the exception of *Gfrα3* and *Ret,* all other genes constitute novel Hoxc8 targets.

We next sought to assess any potential behavioral defects in adult mice lacking *Hoxc8* gene activity in brachial MNs. Due to the perinatal lethality (**Supplementary File 4**), MN axon guidance defects (Catela et al., 2016) and terminal differentiation defects (this study) found in *Hoxc8 MNΔ^early^* mice, we used the *Hoxc8 MNΔ^late^* animals for behavioral analysis. These animals are present in mendelian ratios by weaning stage (**Supplementary File 4**), and *Hoxc8* inactivation in MNs occurs later in development, thereby bypassing the axon guidance defects observed in *Hoxc8 MNΔ^early^* mice. Adult (3 month-old) *Hoxc8 MNΔ^late^* and control *(Hoxc8^fl/fl^* mice were evaluated for rotarod performance. This test evaluates balance and motor coordination by measuring the time spent on an accelerating rotating rod(Deacon, 2013). We found that *Hoxc8 MNΔ^late^* mice tended to perform worse than controls (**Supplementary Fig. 2**). In addition, we conducted a forelimb grip strength test because forelimb muscles are innervated by brachial MNs(Takeshita et al., 2017). Again, *Hoxc8 MNΔ ^late^* mice tended to generate a smaller forelimb muscle force compared to controls (**Supplementary Fig. 2**). These behavioral observations complement our molecular analysis of *Hoxc8 MNΔ^late^* mice, raising the possibility that the observed MN terminal differentiation defects could account for the tendency of *Hoxc8 MNΔ^late^* mice to perform worse than controls.

### In brachial MNs, Hoxc8 partially modifies the suite of its target genes across different life stages

In the context of *C. elegans* MNs, our previous work revealed “temporal modularity” in TF function (Li et al., 2020). That is, the suite of target genes of a continuously expressed TF, in the same cell type, is partially modified across different life stages. Here, we provide evidence for temporal modularity in *Hoxc8* function. We found that the terminal differentiation gene coding for the *glycine receptor subunit alpha-2 (Glra2)*(Young-Pearse et al., 2006) is affected in brachial MNs of *Hoxc8 MNΔ^late^* mice at p8 (**Fig. 4C**). No effect was observed in MNs of *Hoxc8 MNΔ^early^* mice at e12 (**Fig. 4E**), indicating a selective *Hoxc8* requirement for maintenance of *Glra2.* Conversely, the expression of *Ret*, a known *Hoxc8* target gene involved in MN axon guidance(Bonanomi et al., 2012), is selectively reduced in brachial MNs of *Hoxc8 MNΔ^early^* animals at e12 (Catela et al., 2016), but remains unaffected in *Hoxc8 MNΔ ^late^* animals at p8 (**Fig. 4E**), suggesting Hoxc8 is only required for early *Ret* expression. Lastly, Hoxc8 can only activate *Sema5a* expression at embryonic stages (**Fig. 3F-G, Table 1**). Contrary to these stage-specific Hoxc8 dependencies (Hoxc8 controls *Ret* and *Sema5a* at e12 and *Glra2* at p8), we also found that Hoxc8 is continuously required (at e12 and p8) for the expression several terminal differentiation genes *(Nrg1, Mcam, Pappa)* (**Fig. 3F, 4D**). Altogether, these findings suggest that, in brachial MNs, Hoxc8 modifies the suite of its target genes at different developmental stages (**Fig. 4F**). In Discussion, we elaborate on the functional significance of this phenomenon (temporal modularity).

### The Iroquois homeobox (Irx) family of transcription factors constitute ancient Hox targets in motor neurons

In mice, the actions of Hox proteins in motor neuron development are thought to be indirect through intermediary TFs(Philippidou et al., 2012). We therefore wondered whether Hoxc8 controls the expression of terminal differentiation genes via other TFs. Upon interrogating our RNA-Seq data for TFs with reduced expression in brachial MNs of *Hoxc8 MNΔ^eary^* animals (**Fig. 3D-E, Suppl. File 3**), we decided to focus on three members of the Irx family *(Irx2, Irx5, Irx6)* because our analysis in Figure 2 showed that, similar to *Hoxc8,* these TFs are expressed continuously in embryonic and early postnatal brachial MNs (**Fig. 2B**).

Through RNA ISH, we found that expression of *Irx2, Irx5* and *Irx6* is reduced in brachial MNs of *Hoxc8 MNΔ^early^* mice at e12 (**Fig. 5A**), validating the RNA-Seq data. Interestingly, *Irx2* expression is unaffected in MNs of *Hoxc8 MNΔ^late^* mice at p8 (**Fig. 5B**), further supporting the notion of temporal modularity. However, expression of *Irx5* and *Irx6* is reduced in MNs of Hoxc8 *MNΔ^late^* mice at p8 (**Fig. 5B**), suggesting Hoxc8 – similar to its role on the regulation of terminal differentiation genes *(Mcam, Pappa, Nrg1)* – is required for the establishment and maintenance of *Irx5* and *Irx6* expression.

**Figure 5.**
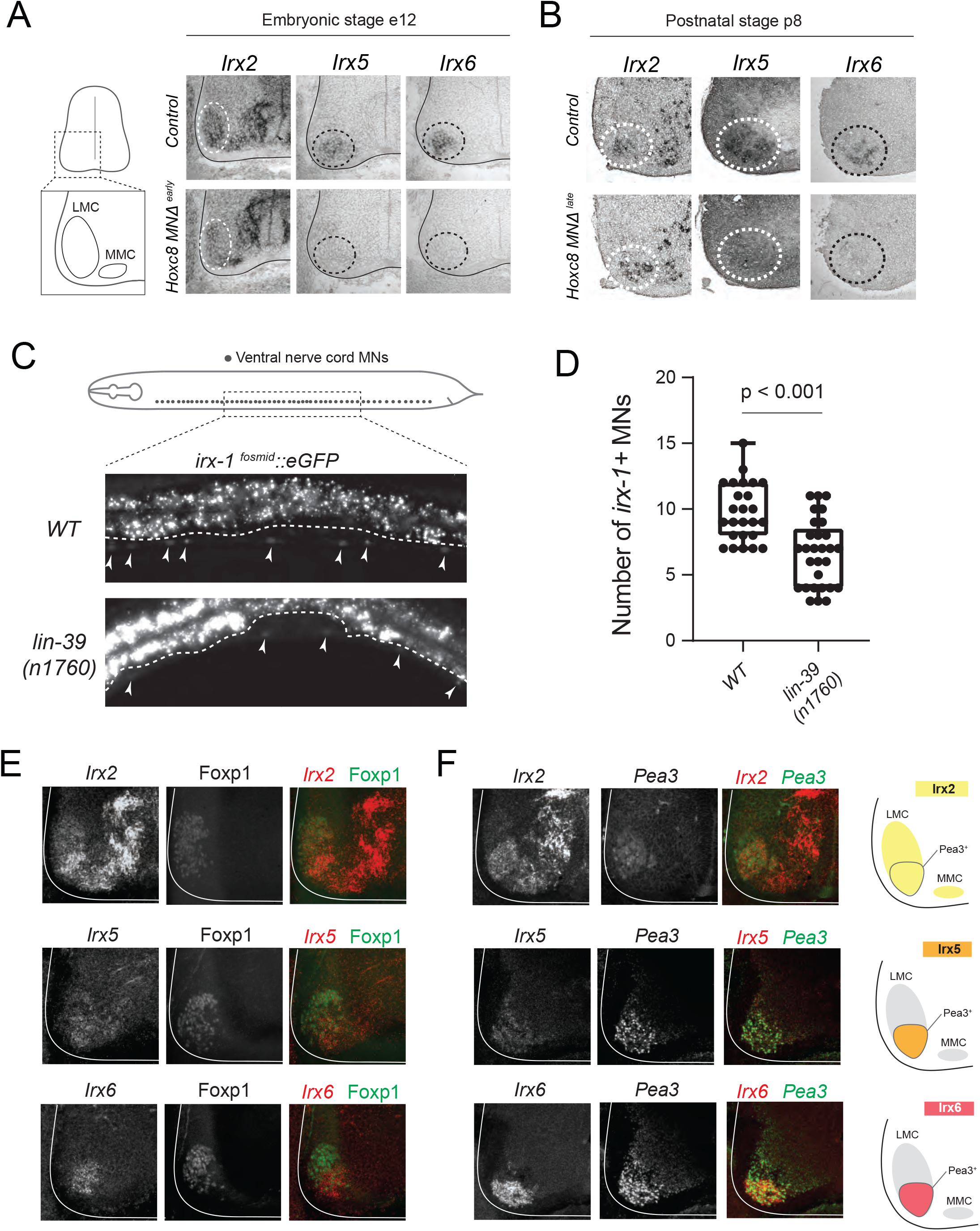
Irx transcription factors are novel Hoxc8 targets in brachial motor neurons. (A) RNA ISH shows reduced *Irx2, Irx5* and *Irx6* expression in e12 *Hoxc8 MNΔ^early^* spinal cords. (B) At p8, *Irx5* and *Irx6* expression is reduced in *Hoxc8 MNΔ^late^* spinal cords, but *Irx2* expression is unaffected (N=4). (C) Expression of a fosmid-based *irx-1* reporter *(wgIs536)* is decreased in C. elegans animals carrying a null *lin-39 (n1760)* allele. A 300μm region of the ventral nerve cord (VNC) was analyzed at the fourth larval stage (L4). Anterior is left, dorsal is up. Arrowheads point to MN nuclei with *irx-1* expression. Green fluorescence signal is shown in white for better contrast. (D) Quantification of total number of MNs in the VNC expressing *irx-1* in WT and *lin-39* null mutants. Box and whiskers plot show the min, max and quartiles with single data points annotated. N > 23. (E – F) RNA ISH for *Irx2, Irx5* and *Irx6* mRNA combined with immunohistochemistry for Foxp1 (LMC marker) and Pea3 (Pea3 pool marker) shows that *Irx2, Irx5* and *Irx6* (in red) are expressed in Foxp1^+^ MNs (green in panel E) and Pea3+ MNs (green in panel F). Red and green signals are shown in white for better contrast. The high levels of *Irx2, Irx5* and *Irx6* expression co-localize with the Pea3 marker. (G) Diagram illustrating the expression pattern of *Irx2, Irx5* and *Irx6* in brachial MNs. Yellow, orange, and red colors indicate expression of *Irx2, Irx5* and *Irx6,* respectively. Gray color indicates no expression.

Our findings on mouse *Hoxc8* (**Fig. 3, 4**) together with previous *C. elegans* studies on *lin-39/Hox* (Feng et al., 2020, Kratsios et al., 2017) strongly suggest an evolutionarily conserved role for Hox in the regulation of MN terminal differentiation genes. Interestingly, we made similar observations regarding the regulation of *Irx* genes. The expression of *irx-1,* the sole *C. elegans* ortholog of the *Irx* family(Petersen et al., 2011), is significantly reduced in nerve cord MNs of nematodes lacking *lin-39/Hox* gene activity (**Fig. 5C-D**). Taken together, these results significantly expand the known repertoire of Hox target genes in the nervous system (**Fig. 4F**), suggesting an ancient role for Hox in the regulation of genes encoding MN terminal differentiation determinants and Irx-type TFs.

### *Irx2* is required for limb-innervating motor neuron development

To test whether Irx proteins play a role in *Hoxc8*-dependent terminal differentiation in mice, we first sought to determine the precise expression pattern of *Irx2, Irx5,* and *Irx6* in subpopulations of brachial MNs. By using specific markers, we found that *Irx2* is expressed in all *Hoxc8+* MNs (LMC and MMC), whereas *Irx5* and *Irx6* are co-expressed with *Hoxc8* in a subgroup of LMC neurons, namely the Pea3 pool (**Fig. 5E-F, Suppl. Figure 3**). Due to its expression in all brachial MNs (**Fig. 5E-G**), we wondered whether *Irx2* controls expression of *Hoxc8*-dependent terminal differentiation genes. We therefore employed CRISPR/Cas9 genome editing and generated *Irx2* mutant mice carrying a 5bp-long deletion in the second exon *(Irx2 Δ^5bp^),* resulting in a premature termination codon (**Fig. 6A-B**) (see Materials and Methods). Mice homozygous for the *Irx2 Δ^5bp^* mutation are viable and fertile. However, we found that the expression of several *Hoxc8*-dependent terminal differentiation genes *(Pappa, Sema5a, Nrg1)* is unaffected in brachial MNs of *Irx2 Δ^5bp^* mutants at e12 (**Suppl. Fig. 4**).

**Figure 6.**
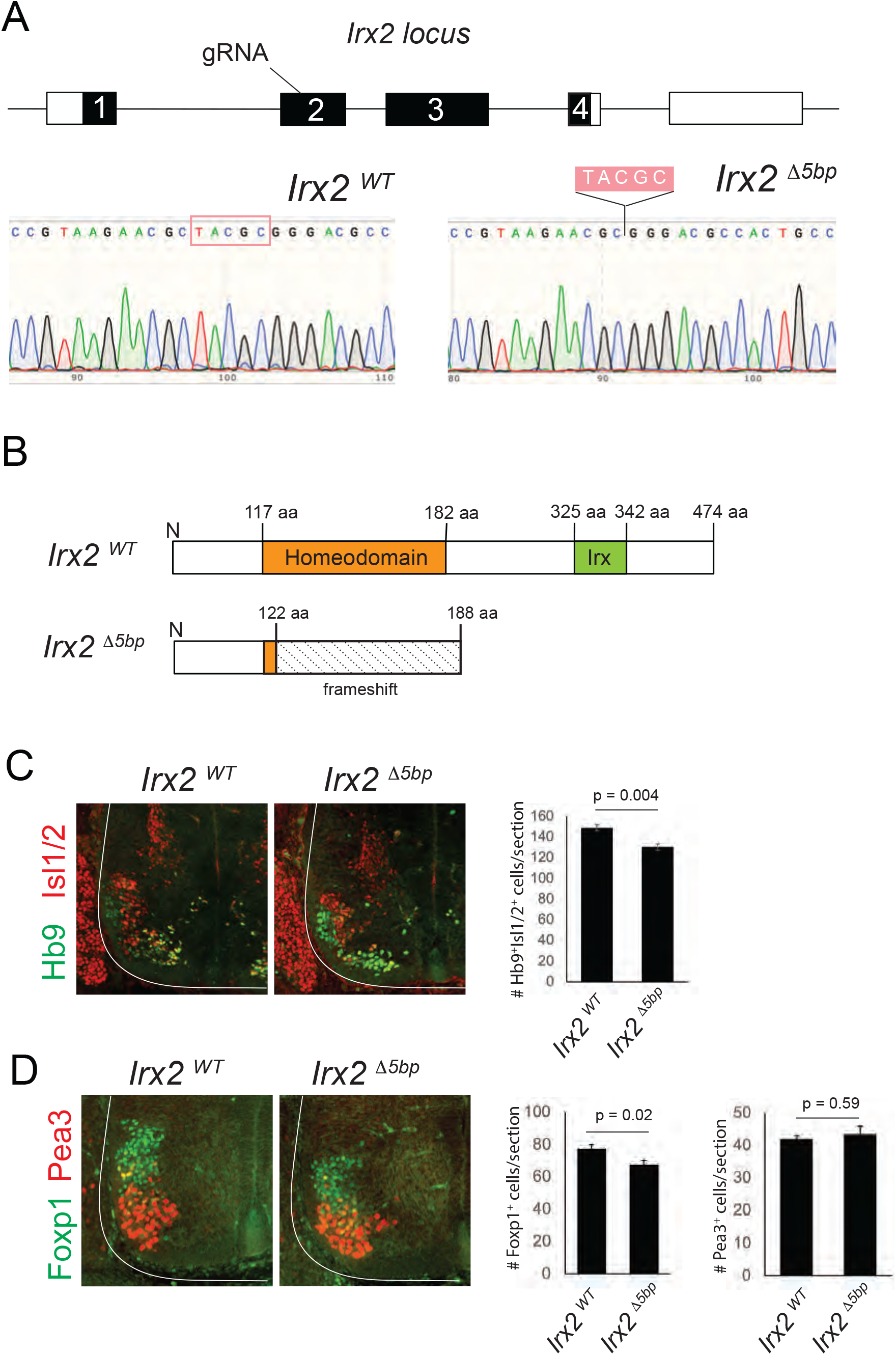
*Irx2* is required for the generation of a subset of limb-innervating MNs. (A) Diagram summarizing the generation of *Irx2*^Δ5bp^ mice. A gRNA was designed to target the second *Irx2* exon, resulting in a CRISPR/Cas9-mediated deletion of 5 base pairs (TACGC) shown in the chromatogram. (B) The *Irx2*^Δ5bp^ allele generates a frameshift mutation and introduces a premature STOP codon resulting in a truncated protein (188 aa instead of 474 aa) that lacks an intact homeodomain and the Irx DNA binding domain. (C-D) Immunostaining for Hb9 and Isl1/2 (panel C), as well as Foxp1 and Pea3 (panel D) shows that the cell body position of MNs is comparable between control and *Irx2^ΔΔ^* spinal cords at e12 (N = 3). However, quantification of Hb9^+^/Isl1/2^+^ motor neurons revealed a significant decrease in total number of brachial motor neurons in *Irx2^ΔΔ^* animals. This decrease is associated with Foxp1 + MNs, and not Pea3^+^ MNs.

We next wondered whether the previously described axonal projection defects of brachial MNs in *Hoxc8 MNΔ^eary^* mice can be explained, at least partially, through the activity of *Irx2.* However, MN axonal projections appeared grossly normal in *Irx2 Δ^5bp^* animals (**Suppl. Fig. 4**). Nevertheless, our analysis uncovered a Hoxc8-independent role for *Irx2* in spinal cord development. We found, at e12, a significant reduction in the total number of brachial MNs (cells co-expressing Hb9 and Isl1/2) in *Irx2 Δ^5bp^* mice (**Fig. 6C**). This reduction occurs in limb-innervating MNs (Devor et al.) located outside the Pea3 pool (Foxp1+ Pea3-) (**Fig. 6D**). Although we cannot pinpoint whether *Irx2* acts at the level of progenitor or post-mitotic cells, the observed reduction in Foxp1+ Pea3-MNs suggests a selective requirement for *Irx2* in the generation and/or survival of limb-innervating MNs.

Hox gene expression is maintained in thoracic and lumbar MNs at postnatal stages In brachial MNs, we found that the expression of multiple Hox genes *(Hoxc4, Hoxa5, Hoxc5, Hoxa6, Hoxc6, Hoxa7, Hoxc8)* is maintained from embryonic to early postnatal stages (Fig. 2). We wondered whether sustained Hox gene expression in MNs is a broadly applicable theme to other rostrocaudal domains of the spinal cord. We therefore performed RNA-Seq on thoracic and lumbar FACS-isolated MNs from *ChAT::IRES::Cre; Ai9* mice at p8 (see Materials and Methods) (Suppl. Fig. 5). Our analysis indeed revealed that, similar to our observations in the brachial domain, Hox genes are expressed postnatally (p8) in thoracic *(Hoxd9)* and lumbar *(Hoxa10, Hoxc10, Hoxa11)* MNs (Suppl. Fig. 5A-C). We further confirmed these findings with RNA ISH (Suppl. Fig. 5D). While the functions of some of these Hox genes are known during the early steps of MN development(Philippidou and Dasen, 2013), their continuous expression suggests additional roles at later embryonic and postnatal stages. Genetic inactivation of these genes at successive developmental stages will determine whether they function in a manner similar to *Hoxc8,* suggesting a more general Hox-based strategy for the control of spinal MN terminal differentiation.

## DISCUSSION

Somatic MNs in the spinal cord innervate hundreds of skeletal muscles and control a variety of motor behaviors, such as locomotion, skilled movement, and postural control. Although we are beginning to understand the molecular programs that control the early steps of spinal MN development(Osseward and Pfaff, 2019, Philippidou and Dasen, 2013, Stifani, 2014), how these clinically relevant cells acquire and maintain their terminal differentiation features (e.g., neurotransmitter phenotype, electrical and signaling properties) remains poorly understood. In this study, we focused on the brachial region of the mouse spinal cord and determined the molecular profile of post-mitotic MNs at a developmental and a postnatal stage. This longitudinal approach identified genes with continuous expression in brachial MNs, encoding novel TFs and effector molecules critical for neuronal terminal differentiation (e.g., ion channels, NT receptors, signaling proteins, adhesion molecules). Interestingly, we also found that most TFs, previously implicated in the early steps of brachial MN development (e.g., initial specification, axon guidance, circuit assembly), such as LIM- and Hox-type TFs (Dalla Torre di Sanguinetto et al., 2008, Philippidou and Dasen, 2013, Stifani, 2014), continue to be expressed in these cells postnatally (p8). Such maintained expression suggested additional roles for these factors during later developmental stages. To test this idea, we focused on *Hoxc8,* identified its target genes, and uncovered a continuous requirement for Hoxc8 in the establishment and maintenance of select MN terminal differentiation features. Our findings dovetail recent Hox studies in the *C. elegans* nervous system (Feng et al., 2020, Kratsios et al., 2017, Zheng et al., 2015) and suggest an evolutionarily conserved role for Hox proteins in the control of neuronal terminal differentiation.

### Expanding the repertoire of Hox target genes in the nervous system

Despite their fundamental roles in patterning the vertebrate hindbrain and spinal cord(Krumlauf, 2016, Parker and Krumlauf, 2017, Philippidou and Dasen, 2013), the downstream targets of Hox proteins in the nervous system remain poorly defined. In this study, we uncovered eight new *Hoxc8* target genes encoding different classes of proteins *(Sema5a* – axon guidance molecule; *Glra2, Nrg1, Mcam, Pappa* – terminal differentiation genes; *Irx2, Irx5, Irx6* – TFs) (**Fig. 4F**), suggesting *Hoxc8* controls different aspects of brachial MN development. While the role of *Hoxc8* in axon guidance and terminal differentiation will be discussed later, we focus here on the identification of *Irx* genes as Hoxc8 targets.

In mice, there are six *Irx* genes *(Irx1-6),* which, similar to Hox genes, are organized into chromosomal clusters (cluster A: *Irx1, Irx2, Irx4;* cluster B: *Irx3, Irx5, Irx6*)(Cavodeassi et al., 2001, Gomez-Skarmeta and Modolell, 2002). During early nervous system development, *Irx* genes play pivotal roles during neuroectoderm specification, forebrain and spinal cord patterning (Briscoe et al., 2000, Cavodeassi et al., 2001, Gomez-Skarmeta and Modolell, 2002, Kobayashi et al., 2002, Houweling et al., 2001, Mummenhoff et al., 2001). At later stages, *Irx* gene function has been primarily studied in the retina, where *Irx4* controls axon guidance, and *Irx5* and *Irx6* are necessary for interneuron terminal differentiation(Cheng et al., 2005, Jin et al., 2003, Star et al., 2012). However, the expression and function of *Irx* genes in the spinal cord remain poorly defined. We found five *Irx* genes to be continuously expressed in post-mitotic spinal MNs, and uncovered a requirement for *Irx2* in the generation and/or survival of limb-innervating MNs. Future gene inactivation studies are needed to determine whether *Irx* genes, similar to their role in the retina, control terminal differentiation features of spinal MNs.

Lastly, our findings shed light into the largely unknown mechanisms that control *Irx* gene expression in the nervous system by identifying *Irx2, Irx5,* and *Irx6* as *Hoxc8* targets. Intriguingly, the sole *C. elegans* ortholog of the *Irx* family *(irx-1)* is also regulated by the Hox protein LIN-39, suggesting *Irx* genes constitute ancient Hox targets in MNs. In *C. elegans, irx-1* is expressed in a subset of ventral nerve cord MNs and controls their identity and connectivity (Petersen et al., 2011). Mouse *Irx5* and *Irx6* may function in an analogous manner, as they are also expressed in a subset of brachial MNs.

### Hoxc8 partially modifies the suite of its target genes to control multiple aspects of brachial MN development

In mice, Hoxc8 is expressed in MNs of the MMC and LMC columns between segments C6 and T1 of the spinal cord(Catela et al., 2016, Tiret et al., 1998), herein referred to as “brachial MNs”. Previous studies using either global *Hoxc8* knock-out or *Hoxc8 MNΔ^eafly^* mice reported aberrant connectivity of forelimb muscles (Catela et al., 2016, Tiret et al., 1998). It was proposed that this early developmental phenotype likely arises due to reduced expression of axon guidance molecules, such as *Ret* and *Gfrα3,* in brachial MNs of *Hoxc8 MNΔ^early^* mice (Catela et al., 2016). Another early developmental defect previously observed in *Hoxc8* mutant mice is the reduced expression of MN pool-specific markers (Scip, Pea3) within the LMC, albeit the overall organization of brachial MNs into MMC and LMC columns appears normal(Catela et al., 2016). Although these findings implicate Hoxc8 in the early steps of brachial MN development, it remained unclear whether Hoxc8 controls additional aspects of MN development during later stages.

In this study, we propose that Hoxc8 controls select features of brachial MN terminal differentiation, such as the expression of the glycine receptor subunit *Glra2,* the cell adhesion molecule *Mcam,* the secreted signaling protein *Pappa,* and a molecule associated with neurotransmission and neuromuscular synapse maintenance *(Nrg1).* We found that all these molecules are expressed continuously in embryonic and postnatal (p8) brachial MNs. By removing Hoxc8 gene activity either at an early *(Hoxc8 MNΔ^eafly^* mice) or late *(Hoxc8 MNΔ^late^* mice) developmental stage, we uncovered a continuous Hoxc8 requirement for the initial expression and maintenance of *Mcam, Pappa,* and *Nrg1.* Intriguingly, we also found evidence for temporal modularity in Hoxc8 function, that is, the suite of Hoxc8 targets in brachial MNs is partially modified at different developmental stages. Three line of evidence support this notion: (a) expression of the terminal differentiation gene *Glra2* is only affected in *Hoxc8 MNΔ^late^* mice, indicating a selective *Hoxc8* requirement for *Glra2* maintenance in MNs, (b) expression of two axon guidance molecule *(Sema5a, Ret)* is only affected in MNs of *Hoxc8 MNΔ^early^* mice, and (c) *Irx2* expression is only affected in *Hoxc8 MNΔ^early^* mice.

What is the purpose of such temporal modularity? We propose that Hoxc8 partially modifies the suite of its target genes at different life stages in order to control different facets of brachial MN development, such as axon guidance and terminal differentiation (**Fig. 4F**). During early development, Hoxc8 controls axon guidance molecules, such as *Ret* (Bonanomi et al., 2012, Catela et al., 2016) and *Sema5a* (this study) in order to ensure proper MN-muscle connectivity. Consistent with this idea, similar axon guidance defects occur in *Hoxc8* and *Ret* mutant mice(Catela et al., 2016). During late development, Hoxc8 maintains the expression of the glycine receptor subunit *Glra2,* a terminal differentiation marker necessary for glycinergic input to brachial MNs(Young-Pearse et al., 2006). Apart from Hoxc8, temporal modularity has been recently described for two other TFs: UNC-3 in *C. elegans* MNs and Pet-1 in mouse serotonergic neurons(Li et al., 2020, Wyler et al., 2016). Like Hoxc8, UNC-3 and Pet-1 control various aspects of *C. elegans* motor and mouse serotonergic neurons (e.g., axon guidance, terminal differentiation)(Donovan et al., 2019, Kratsios et al., 2012, Liu et al., 2010, Prasad et al., 1998). Although the mechanistic basis of such modularity remains poorly understood, a possible scenario is the employment of transient enhancers – a mechanism recently proposed for maintenance of gene expression in *in vitro* differentiated spinal MNs (Rhee et al., 2016). We surmise that temporal modularity in TF function may be a broadly applicable mechanism enabling a single TF to control different, temporally segregated “tasks/processes” within the same neuron type.

### A new role for Hox in the mouse nervous system: Establishment and maintenance of neuronal terminal differentiation

Much of our current understanding of Hox protein function in the nervous system stems from studies in the vertebrate hindbrain and spinal cord, as well as the *Drosophila* ventral nerve cord(Baek et al., 2013, Baek et al., 2019, Estacio-Gomez and Diaz-Benjumea, 2014, Estacio-Gomez et al., 2013, Karlsson et al., 2010, Mendelsohn et al., 2017, Miguel-Aliaga and Thor, 2004, Moris-Sanz et al., 2015, Parker and Krumlauf, 2017, Philippidou and Dasen, 2013). This large body of work has established Hox proteins as critical regulators of the early steps of neuronal development including cell specification, migration, survival, axonal path finding, and circuit assembly. However, the functions of Hox proteins in later steps of nervous system development remain poorly understood. Recent work on invertebrate Hox genes has begun to address this knowledge gap. In *Drosophila* MNs necessary for feeding, *Deformed (Dfd)* is required to maintain neuromuscular synapses(Friedrich et al., 2016). In *C. elegans* touch receptor neurons, the anterior *(ceh-13)* and posterior *(egl-5)* Hox genes control the expression levels of the LIM homeodomain protein MEC-3, which in turn controls touch receptor terminal differentiation (Zheng et al., 2015). In the *C. elegans* ventral nerve cord, midbody *(lin-39, mab-5)* and posterior *(egl-5)* Hox genes control distinct terminal differentiation features of midbody and posterior MNs, respectively(Kratsios et al., 2017). LIN-39 binds to the *cis*-regulatory region of multiple terminal differentiation genes (e.g., ion channels, NT receptors, signaling molecules) and is required for their maintained expression in MNs during post-embryonic stages(Feng et al., 2020).

Our Hoxc8 findings in mice support the hypothesis that Hox-mediated control of later aspects of neuronal development (e.g. terminal differentiation) is evolutionarily conserved from invertebrates to mammals. Similar to *C. elegans* Hox genes, mouse *Hoxc8* is continuously expressed in brachial MNs from embryonic to early postnatal stages, and sustained *Hoxc8* gene activity is required to establish and maintain at later developmental stages the expression of several terminal differentiation genes. This noncanonical, late function of Hoxc8 may be shared by other Hox genes in the mouse nervous system. In the spinal cord, we found that several Hox genes are expressed in brachial *(Hoxc4, Hoxa5, Hoxc5, Hoxa6, Hoxc6, Hoxa7),* thoracic *(Hoxd9)* and lumbar *(Hoxc10, Hoxa11)* MNs at postnatal (p8) stages. Moreover, a recent study reported expression of 24 Hox genes in the adult mouse brainstem, and hypothesized that maintained Hox gene expression is necessary for activity-dependent synaptic pruning and maturation(Hutlet et al., 2016). To date, the functional significance of maintained Hox gene expression in the CNS remains largely unknown, and temporally controlled genetic approaches are required to fully elucidate the late functions of this remarkable class of highly conserved TFs.

### The quest for terminal selectors of spinal motor neuron identity

Numerous genetic studies in the nematode *C. elegans* support the idea that continuously expressed TFs (termed “terminal selectors”) establish during development and maintain throughout postembryonic life the identity and function of individual neuron types by activating the expression of terminal differentiation genes (e.g., NT biosynthesis components, ion channels, adhesion and signaling molecules)(Deneris and Hobert, 2014, Hobert, 2008, Hobert, 2016). Multiple cases of terminal selectors for various neuron types have already been described in flies, cnidarians, marine chordates, and mice, suggesting deep conservation for this type of regulators(Allan and Thor, 2015, Deneris and Hobert, 2014, Hobert, 2008, Hobert, 2016, Hobert and Kratsios, 2019, Tourniere et al., 2020). However, it remains unclear whether spinal MNs in vertebrates employ a terminal selector-type of mechanism to acquire and maintain their terminal differentiation features. Addressing this knowledge gap could aid the development of *in vitro* protocols for the generation of mature and terminally differentiated spinal MNs, a much anticipated goal in the field of MN disease modeling(Sances et al., 2016).

Two lines of evidence implicate *Hoxc8* in the control of MN terminal differentiation: (a) *Hoxc8* is expressed continuously, from embryonic to early postnatal stages, in brachial MNs, and (b) both early and late removal of *Hoxc8* in brachial MNs affected the expression of several terminal differentiation genes, suggesting continuous requirement. However, *Hoxc8* does not act alone – loss of *Hoxc8* did not completely eliminate the expression of its target genes (Fig. 3F, 4C). This residual expression indicates that additional TFs are necessary to control brachial MN terminal differentiation. One such factor is the LIM homeodomain protein *Islet1 (Isl1),* which is required for early induction of genes necessary for ACh biosynthesis in mouse spinal MNs and the *in vitro* generation of MNs from human pluripotent stem cells(Cho et al., 2014, Qu et al., 2014, Rhee et al., 2016). Interestingly, *Isl1* is expressed continuously in brachial MNs (Fig. 2) and amplifies its own expression (Erb et al., 2017) – both defining features of a terminal selector gene. In addition to *Isl1,* our expression analysis revealed multiple TFs from different families (e.g., Hox, Irx, LIM) with continuous expression in brachial MNs (Fig. 2, Table 2). In the future, temporally controlled gene inactivation studies are needed to determine whether these TFs participate in the control of spinal MN terminal differentiation. Intriguingly, the majority of the TFs with continuous expression in brachial MNs belong to the homeodomain family. Homeodomain TFs are overrepresented in the current list of *C. elegans* and mouse terminal selectors(Deneris and Hobert, 2014, Reilly et al., 2020, Serrano-Saiz et al., 2013), suggesting an ancient role for this family of regulatory factors in the control of neuronal terminal differentiation.

## ACKNOWLEDGEMENTS

We are grateful to members of the Kratsios lab (Yinan Li, Edgar Correa, Nidhi Sharma, Filipe Goncalves Marques) and Drs. Deeptha Vasudevan, Ellie Heckscher, and Oliver Hobert for comments on the manuscript. We thank Dr. Jeremy Dasen (NYU) for providing the following antibodies (rabbit anti-Foxp1, rabbit anti-Lhx3, rabbit anti-Hb9, rabbit anti-Isl1/2, rabbit anti-Pea3) and Jihad Aburas for technical assistance. We thank the following Core Facilities at The University of Chicago: (a) Cytometry and Antibody Technology, (b) Genomics Facility (RRID:SCR_019196), especially Dr. Pieter Faber, for his assistance with the RNA-Sequencing, and (c) Transgenic Mouse and Embryonic Stem Cell Facility, especially Linda Degenstein, for the generation of Irx2 mutant mice using CRISPR/Cas9 genome editing. This work was supported by the Lohengrin Foundation (P.K.) and a grant from the National Institute of Neurological Disorders and Stroke (NINDS) of the NIH (Award Number: R01NS116365) to P.K.

## AUTHOR CONTRIBUTIONS

C. C., Conceptualization, Data curation, Investigation, Visualization, Methodology, Writing – original draft, review and editing; Y.W., K. *W., W.F.,* Formal analysis, Validation, Investigation; P. K., Conceptualization, Supervision, Investigation, Funding acquisition, Project administration, Writing— original draft, review and editing.

## DECLARATION OF INTERESTS

The authors declare no competing interests.

## DATA ACCESSIBILITY

Sequencing data have been deposited in GEO under accession code GSE174709. All data generated or analyzed in this study are included in the manuscript and supporting files.

The following GEO dataset was generated:

Authors: Catarina Catela, Paschalis Kratsios

Year: 2021

Dataset Title: New roles for Hoxc8 in the establishment and maintenance of motor neuron identity

Dataset URL: https://www.ncbi.nlm.nih.gov/geo/query/acc.cgi?acc=GSE174709

Database and Identifier: NCBI Gene Expression Omnibus, GSE174709

## MATERIALS AND METHODS

### Mouse husbandry and genetics

All mouse procedures were approved by the Institutional Animal Care and Use Committee (IACUC) of the University of Chicago. Generation of conditional *Hoxc8*(Blackburn et al., 2009), *Olig2::Cre* (Dessaud et al., 2007), *Hb9::GFP* (Wichterle et al., 2002), *ChAT-IRES-Cre* (Rossi et al., 2011) and *Ai9* (Madisen et al., 2010) mice have been previously described. The *Irx2* mutant mice were generated at the Trangenics/ES Cell Technology Mouse Core Facility of the University of Chicago. A gRNA targeting exon 2 of *Irx2* (gRNA: GTACCGTAAGAACGCTACGC) was injected into mouse zygotes together with a Cas9-expressing plasmid. Founder animals were genotyped by sequencing and a 5 bp deletion was detected in exon 2 of the *Irx2* locus *(Irx2 Δ^5bp^).* In the F1 generation, heterozygous *Irx2 Δ^5bp^* mice were used for crosses to establish a homozygous mutant line. Primers for genotyping the *Irx2 Δ^5bp^* allele: Irx2 (5’-ACAGGCTGTTGTGGGTTCC-3’ and 5’-CATCCTGTGCCTTGTCTGAA-3’).

### Motor neuron fluorescence activated cell sorting (FACS) and RNA extraction

For the analysis shown in Figure 1, brachial MNs from e12.5 *Hb9::GFP* and p8 *ChAT-IRES-Cre; Ai9* animals were isolated from segments C4-T1 using the dorsal root ganglia as reference. For the analysis shown in Figure 3D, segments C7-T2 were used. Neurons were dissociated using papain and filtered (using 50μm filters) for sorting. A GFP negative spinal cord was also included as a negative control for the FACS setup. DAPI staining was used to excluded dead cells from the sorting. Cells were collected into Arcturus Picopure extraction buffer, and immediately processed for RNA isolation. RNA was extracted from purified MNs, using the Arcturus Picopure RNA isolation kit (Arcturus, #KIT0204). For the RNA-Seq analysis on e12.5 *Hb9::GFP* embryos, three biological replicates were used; 5-6 spinal cords were pooled per replicate. For the RNA-Seq analysis on p8 *ChAT-IRES-Cre; Ai9* animals, three biological replicates were used; 3 spinal cords were pooled per replicate. RNA quality and quantity were measured with an Agilent Picochip (Agilent 2100 Bioanalyzer). All samples had high quality scores between 9-10 RIN. After cDNA library preparation, RNA sequencing was performed using a 50-nucleotide single-end read rapid run flow cell lanes with the Illumina HiSeq 4000 sequencer (University of Chicago genomics Core facility).

### RNA-Seq analysis

Raw sequence data were subjected to quality control using the FastQC algorithm (http://www.bioinformatics.babraham.ac.uk/projects/fastqc/). Unique reads were aligned into the mouse genome (GRCm38/mm10) using the HISET2 alignment program (Kim et al., 2015) followed by transcript counting with the featureCounts program (Liao et al., 2014). Differential gene expression analysis between embryonic and postnatal samples was performed with the DESeq2 program (Love et al., 2014). All analyses were performed using the open source, web-based Galaxy platform (https://usegalaxy.org). The heatmaps were generated using the Morpheus program developed by the Broad Institute (https://software.broadinstitute.org/morpheus). Gene hierarchical clustering was performed using a Pearson’s correlation calculation.

### RNA *in situ* hybridization

E12.5 embryos and p8 spinal cords were fixed in 4% paraformaldehyde for 1.5–2 h and overnight, respectively, placed in 30% sucrose overnight (4 °C), and embedded in optimal cutting temperature (OCT) compound. Cryosections were generated and processed for *in situ* hybridization or immunohistochemistry as previously described (Dasen et al., 2005, De Marco Garcia and Jessell, 2008). For double fluorescence *in situ* hybridization, two different probes were used, one labeled with digoxigenin and the other with biotin. Anti-DIG-peroxidase conjugated (Roche) and anti-biotin-peroxidase conjugated antibodies were used (Vector Labs). RNA was detected using a fluorescein and a Cy3 Tyramide Amplificafion system (Perkin Elmer). Combination of fluorescent *in situ* hybridization with antibody staining was performed as follows: cryosections were postfixed in 4% paraformaldehyde, washed in PBS, endogenous peroxidase was blocked with a 0.1% H2O2 solution and permeabilized in PBS/0.1% Triton-X100. Upon hybridization with DIG-labeled RNA probe overnight at 72 °C and washes in SSC, the anti-DIG antibody conjugated with peroxidase (Roche) and primary antibody against Pea3 (rabbit anti-Pea3, Dr. Jeremy Dasen) or Foxp1 (rabbit anti-Foxp1, Dr. Jeremy Dasen) were applied overnight (4 °C) to the sections. The next day, the sections were incubated with the secondary antibody (Alexa 488 donkey anti-rabbit IgG, Life Technologies, A21206) and detection of RNA was performed using a Cy3 Tyramide Amplification system (Perkin Elmer). Images were obtained with a high-power fluorescent microscope (Zeiss Imager V2) and analyzed with Fiji software (Schindelin et al., 2012).

### Immunohistochemistry and wholemount GFP immunostaining

Fluorescence staining on cryosections and wholemount GFP immunostaining were performed as previously described (Catela et al., 2016).

### Gene Ontology analysis

Protein classification was performed using the Panther Classification System Version 15.0 (http://www.pantherdb.org). Embryonic (1381 out of 2904) and postnatal (1348 out of 2699) motor neuron genes were categorized into protein classes using the algorithms built into Panther (Mi et al., 2013, Thomas et al., 2003).

### Rotarod performance test

Age-matched (3-month old) female mice were trained on an accelerating rotarod for 5 days. The experimenter was blind to the genotypes. A computer-controlled rotarod apparatus (Rotamex-5, Columbus Instruments, Columbus, OH, USA) with a rat rod (7 cm diameter) was set to accelerate from 4 to 40 revolutions per minute (rpm) over 300 s, and recorded time to fall. Mice received five consecutive trials per session, one session per day (about 30 s between trials).

### Forelimb grip strength test

The forelimb strength of age-matched (3-month old) male mice was measured using a grip strength meter from Bioseb (model BIO-GS3). We followed the manufacturer’s protocol. In brief, the meter was positioned horizontally and mice were held by the tail and lowered towards the apparatus. The mice were allowed to grasp the metal grid and were then pulled backwards in the horizontal plane. The maximum force of grip was measured and we used the average of five measurements for analysis. Force was measured in Newton.

### Quantification of *irx-1* reporter gene expression in *C. elegans* MNs

Worms were grown at 20°C on nematode growth media (NGM) plates seeded with bacteria *(E.coli* OP50) as food source (Brenner, 1974). Animals carrying the *lin-39 (n1760)* mutant allele were crossed with animals carrying the *wgIs536 [irx-1::TY1::EGFP::3xFLAG + unc-119 (+)]*fosmid-based reporter for *irx-1.* Homozygous animals for the *lin-39 (n1760)* allele and the *wgIs536* reporter were anesthetized at the fourth larval stage (L4) using 100mM of sodium azide (NaN_3_) and mounted on a 4% agarose pad on glass slides. Images were taken using an automated fluorescence microscope (Zeiss, Axio Imager Z2). Acquisition of several z-stack images (each ~1 μm thick) was taken with Zeiss Axiocam 503 mono using the ZEN software (Version 2.3.69.1000, Blue edition). Representative images are shown following max-projection of 1-8 μm Z-stacks using the maximum intensity projection type. Image reconstruction was performed using Fiji (Schindelin et al., 2012).

### Statistical analysis

For data quantification, graphs show values expressed as mean ± standard error of the mean (SEM). With the exception of the rotarod experiment, all other statistical analyses were performed using the unpaired *t*-test (two-tailed). Differences with *p* < 0.05 were considered significant. For the rotarod performance test, two-way ANOVA was performed (Prism Software, Genotype effect, F [1,9] = 2.097; P = 0.18; Time X Genotype interaction, F [24, 216] = 1.196, P = 0.24).

## SUPPLEMENTARY FIGURE LEGENDS

**Suppl. Figure 1. RNA ISH analysis of terminal differentiation markers in *Hoxc8 MNΔ^early^* mice**. Expression of *Slc18a3, Slc5a7, Glra2, Slc10a4, Slc44a5, Sncg, Ngfr, Nyap2, Pex14* and *Tagln2* was detected in the ventrolateral region of e12.5 spinal cords in control *(Hoxc8^fl/fl^* and *Hoxc8 MNΔ^early^* embryos (N = 4).

**Suppl. Figure 2. Behavioral analysis of *Hoxc8 MNΔ^late^* mice.** (A) Rotarod performance test showed a trend of lower latency/second in 3 month-old *Hoxc8 MNΔ^late^* female mice (N = 5) compared to control *(Hoxc8 fl/fl)* mice (N = 6). The following statistical analysis was performed: Two-way ANOVA (Prism Software), Genotype effect, F (1,9) =2.097, P=0.18; Time X Genotype interaction, F (24,216) = 1.196, P= 0.24).

(B) Forelimb grip strength was performed on 3 month-old *Hoxc8 MNΔ^late^* male (N = 3) and control *(Hoxc8 fl/fl)* mice (N = 5). Force was measured in Newton.

**Suppl. Figure 3. Expression of *Irx2, Irx5,* and *Irx6* at brachial, thoracic and lumbar domains of the embryonic spinal cord.** RNA ISH showing expression of *Irx2* at the brachial, thoracic and lumbar domains of a wild-type e12.5 spinal cord (N = 4). Expression of *Irx5* and *Irx6* is mainly detected at brachial and lumbar regions. *Irx2, Irx5,* and *Irx6* are expressed in the ventrolateral region, which is populated by MNs. A schematic representation of motor column (MC) location is provided at the top (LMC: lateral MC; MMC: Medial MC, HMC: Hypaxial MC; PGC: Preganglionic MC).

**Suppl. Figure 4. Characterization of brachial motor neurons of *Irx2 Δ^5bp^* mutant mice.**

(A) RNA ISH showing expression of *Nrg1, Sema5a and Pappa* in the e12.5 spinal cord of control and *Irx2 Δ^5bp^* homozygous mutant mice. N = 4. (B) Wholemount GFP immunostaining showing that there are no differences in MN axonal projections to the forelimb and trunk between control and *Irx2 Δ^5bp^* homozygous mutant mice at e12.5. N = 4.

**Suppl. Figure 5. Different Hox genes are expressed in brachial, thoracic, and lumbar MNs at postnatal day 8.**

(A) Schematic showing the different regions of the p8 spinal cord (brachial, thoracic, lumbar) used for RNA-Seq analysis. The *ChAT-IRES-Cre; Ai9* mouse line was used to fluorescently label MNs (see Materials and Methods). (B) MA plots showing the differential expression of genes (each dot is an individual transcript) in brachial, thoracic, and lumbar regions. (C) Plots showing the normalized counts of *Hoxc5, Hoxc8, Hoxc9, Hoxa10* and *Hoxc10* transcripts in the brachial, thoracic, and lumbar regions of the spinal cord. (D) *RNA ISH* was used to independently validate the RNA-Seq data shown in panels B-C. Expression of *Hoxc4, Hoxa5, Hoxc5, Hoxa6, Hoxc6, Hoxa7, Hoxc8* was assessed in p8 brachial MNs, and is also shown in Figure 2 (panel A). Expression of *Hoxd9* was assessed in p8 thoracic MNs, whereas *Hoxc10* and *Hoxa11* were evaluated in p8 lumbar MNs (N = 4).

## Notes

### Competing Interest Statement

The authors have declared no competing interest.

