## Supplementary figures and images for "Control of spinal motor neuron terminal differentiation through sustained *Hoxc8* gene activity"

### Supplementary Figures 1 - 5

#### Suppl. Figure 1

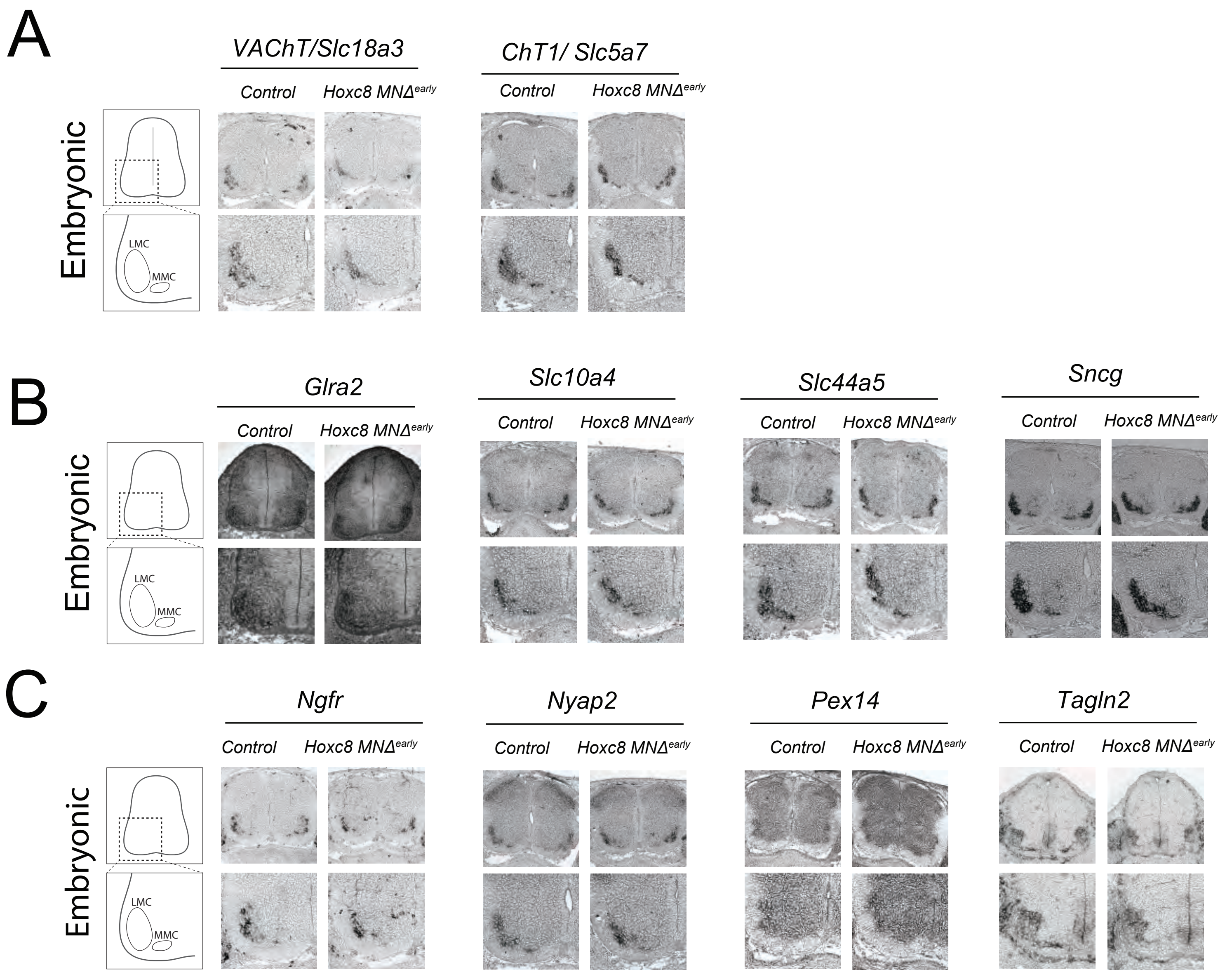

Suppl. Figure 2

A

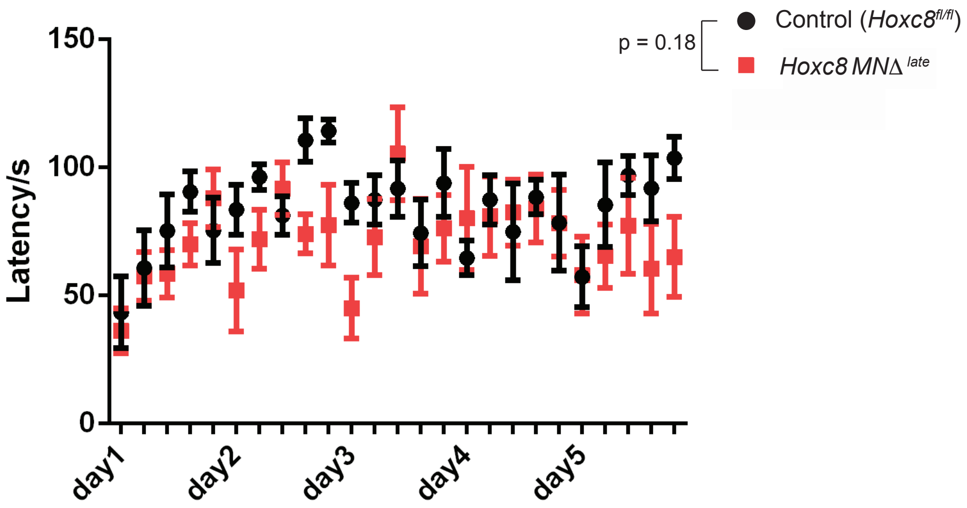


B

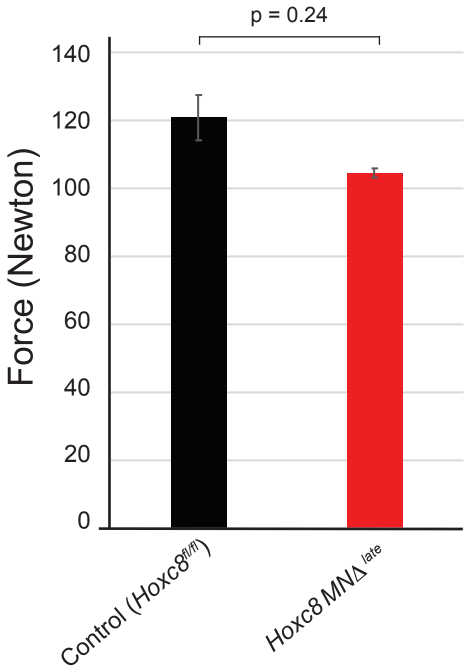

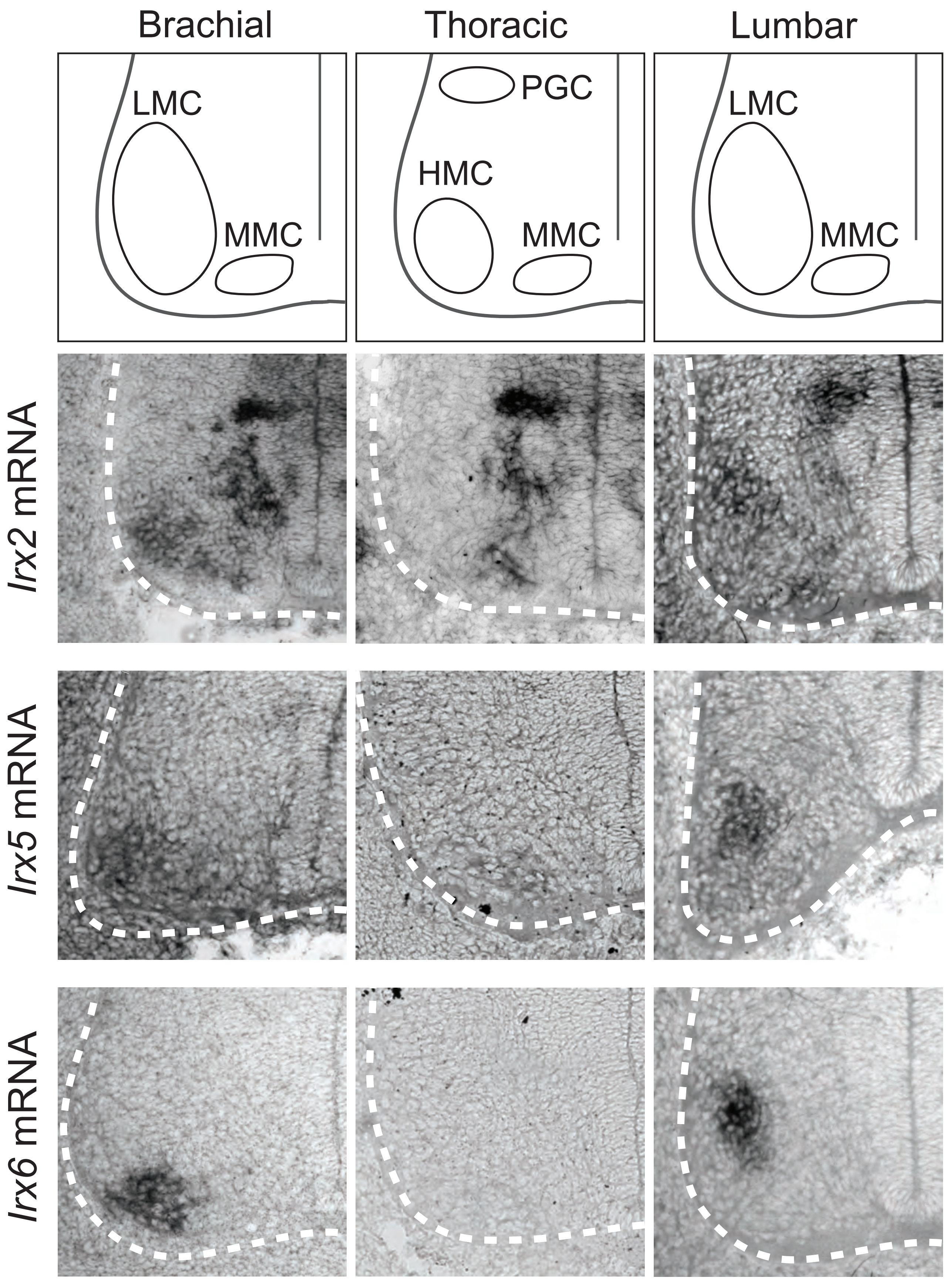

Suppl. Figure 4

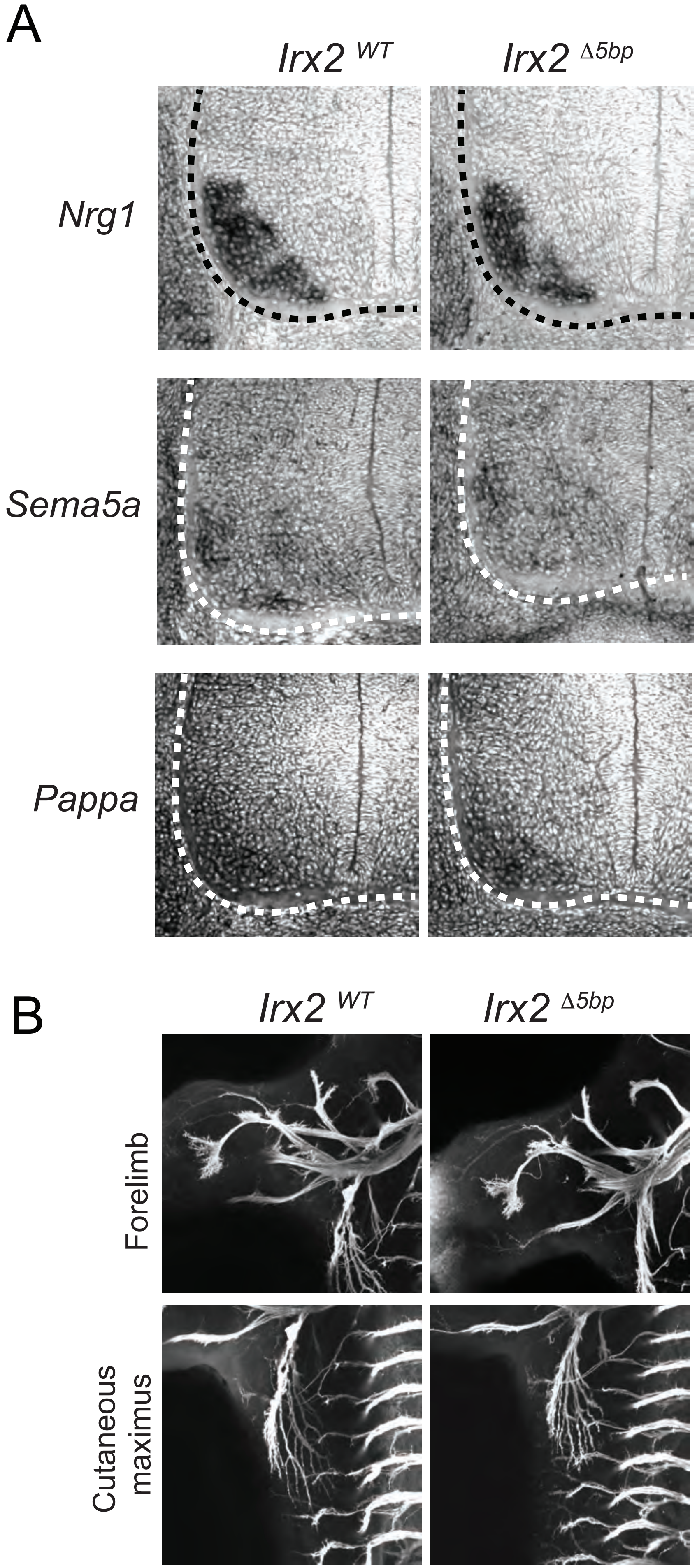

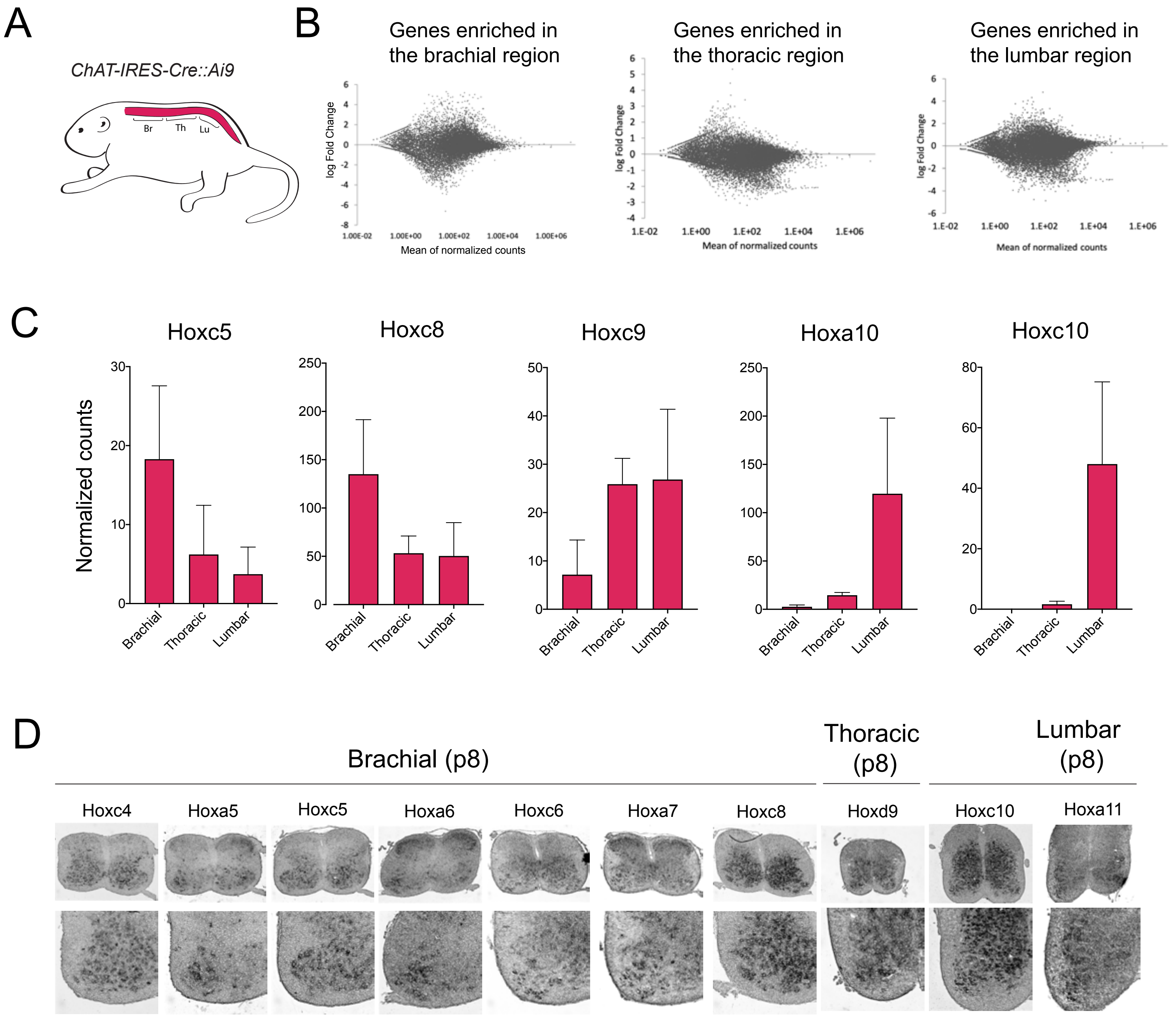
